# Extended regulation interface coupled to the allosteric network and disease mutations in the PP2A-B56δ holoenzyme

**DOI:** 10.1101/2023.03.09.530109

**Authors:** Cheng-Guo Wu, Vijaya K. Balakrishnan, Pankaj S. Parihar, Kirill Konovolov, Yu-Chia Chen, Ronald A Merrill, Hui Wei, Bridget Carragher, Ramya Sundaresan, Qiang Cui, Brian E. Wadzinski, Mark R. Swingle, Alla Musiyenko, Richard Honkanen, Wendy K. Chung, Aussie Suzuki, Stefan Strack, Xuhui Huang, Yongna Xing

## Abstract

An increasing number of mutations associated with devastating human diseases are diagnosed by whole-genome/exon sequencing. Recurrent *de novo* missense mutations have been discovered in B56δ (encoded by *PPP2R5D*), a regulatory subunit of protein phosphatase 2A (PP2A), that cause intellectual disabilities (ID), macrocephaly, Parkinsonism, and a broad range of neurological symptoms. Single-particle cryo-EM structures show that the PP2A-B56δ holoenzyme possesses closed latent and open active forms. In the closed form, the long, disordered arms of B56δ termini fold against each other and the holoenzyme core, establishing dual autoinhibition of the phosphatase active site and the substrate-binding protein groove. The resulting interface spans over 190 Å and harbors unfavorable contacts, activation phosphorylation sites, and nearly all residues with ID-associated mutations. Our studies suggest that this dynamic interface is close to an allosteric network responsive to activation phosphorylation and altered globally by mutations. Furthermore, we found that ID mutations perturb the activation phosphorylation rates, and the severe variants significantly increase the mitotic duration and error rates compared to the wild variant.

## Introduction

Protein phosphatase 2A (PP2A) is a major serine/threonine phosphatase in the PPP family that targets many cellular phosphoproteins via diverse heterotrimeric holoenzymes in mammalian cells^1–6^. Each holoenzyme consists of a common core formed by the scaffolding A and catalytic C (PP2Ac) subunits and a diverse regulatory subunit from one of the four major families (B/B55/PR55 (*PPP2R2*), B’/B56/PR61 (*PPP2R5*), B’’/PR72 ((*PPP2R3*), and B’’’/Striatin).

Whole exome/genome sequencing identified mutations in PP2A subunits in cancer and largely *de novo* mutations in neurological disorders^7–11^. Both unique and recurrent mutations have been found in the common core of the B56 regulatory subunits. Up to 20 recurrent missense mutations in B56δ cause severe intellectual and developmental disorders^8, 9, 12, 13^, known as Jordan Syndrome. More recently, multiple missense variants in B56δ have been associated with early-onset Parkinsonism^14–16^. The incidence of neurodevelopmental and neurodegenerative disorders associated with B56δ is estimated at 2.32 to 2.87 per 100,000 births; 250,000 cases are estimated to be undiagnosed^8, 9, 12, 14–16^.

Recent studies indicate that PP2A holoenzymes from the B56 family recognize a signature short linear motif (SLiM), LxxIxE, in the disordered regions of substrates^17–19^. Diverse SLiMs are found in intrinsically disordered regions that serve as docking interfaces for peptide-binding proteins and play crucial roles in modulating cellular signaling^20, 21^. Several SLiMs have been uncovered for the PPP family phosphatases targeting different substrates^22–25^. The B56-targeting LxxIxE SLiM binds to a protein groove in the conserved common core of B56 regulatory subunits^19, 26^. Several thousands of B56 SLiM-containing proteins are predicted in the human proteome. Many of these are involved in broad cellular and physiological processes^17^. B56 regulatory subunits play essential roles in neurodevelopment, brain functions, and tumor suppression, as reflected by their ability to control cell cycle^17, 18, 27–29^, cytoskeleton dynamics^30^, DNA damage responses^31, 32^, CREB signaling^33^, and c-Myc stability^34–38^.

In addition to the common core, B56δ possesses long disordered regions at the N- and C-termini that harbor multiple phosphorylation sites (Extended Data Figs. 1-2). The PP2A-B56δ holoenzyme is known to be highly regulated by distinct cellular signaling pathways. Protein kinase A (PKA) phosphorylates B56δ and activates the holoenzyme, thereby regulating signaling molecules downstream of cyclic adenosine monophosphate (cAMP)^39–41^. At the G2/M checkpoint, the DNA-responsive checkpoint kinase Chk1 phosphorylates B56δ and stimulates the holoenzyme activity^42^. B56δ also plays a critical role in controlling mitotic exit^43^. Moreover, the B56δ holoenzyme is phosphorylated by the ataxia-telangiectasia mutated (ATM) kinase upon DNA damage and regulates p53 function^44^. Understanding the structure of the PP2A-B56δ holoenzyme and the mechanisms by which it is activated is essential for shedding light on its complex regulation and the pathological mechanisms of its disease mutations.

**Figure 1.**
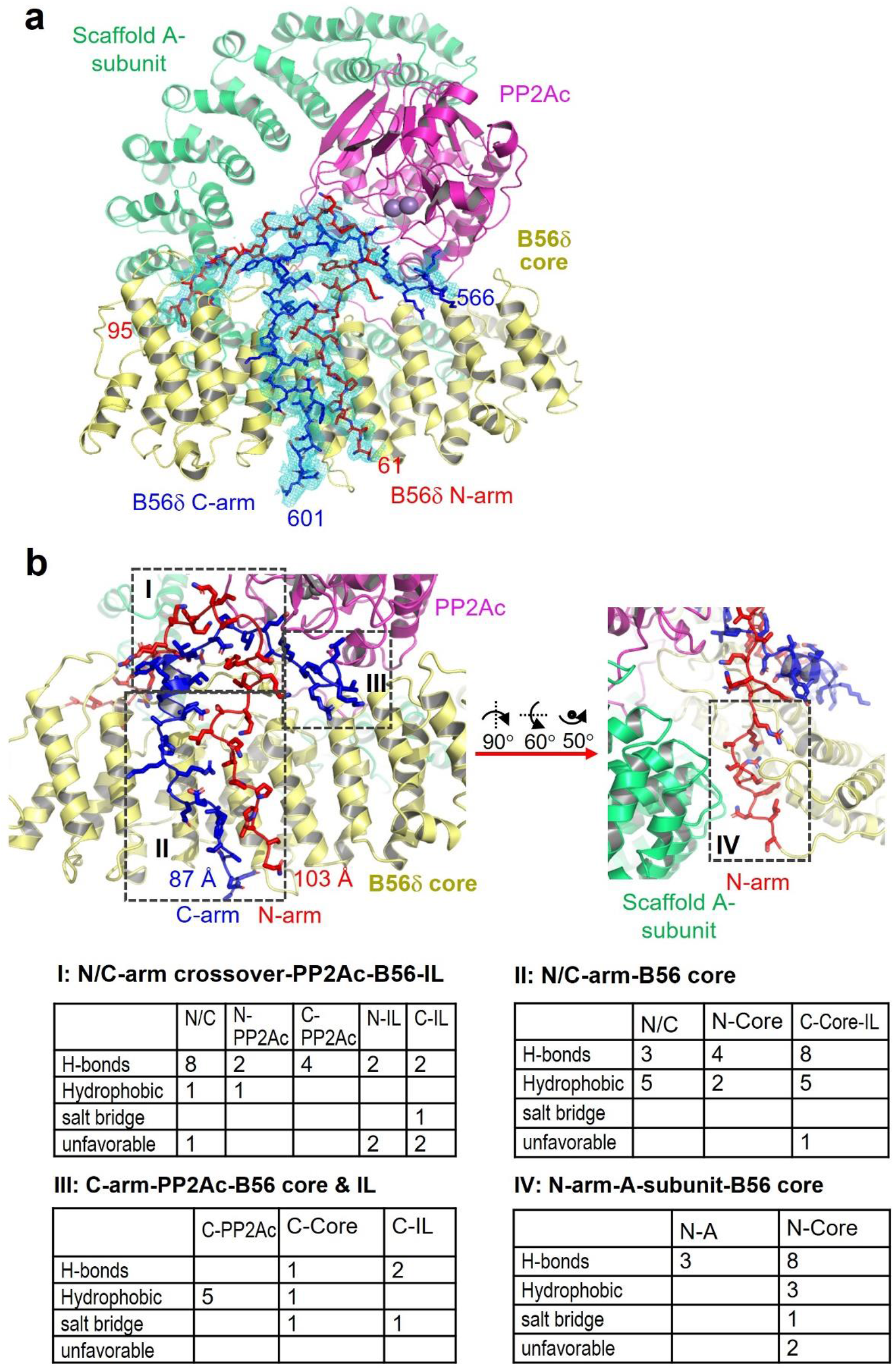
The cryo-EM structure of the closed form of the E197K PP2A-B56δ holoenzyme. (a) The overall cryo-EM structure of the E197K holoenzyme in the closed form. The A subunit, PP2Ac, B56δ core, and N/C-arms are colored green, magenta, yellow, red, and blue, respectively. The electron density map for N/C-arms is colored cyan. The N/C-arms are in sticks and the rest of the structure is in cartoon. Manganese ions are shown in grey spheres. (b) Mapping of the contact properties along the path of the super-dynamic interface spanning over 190 Å. The presentation of the structural model and the color scheme are the same as in (a).

**Figure 2.**
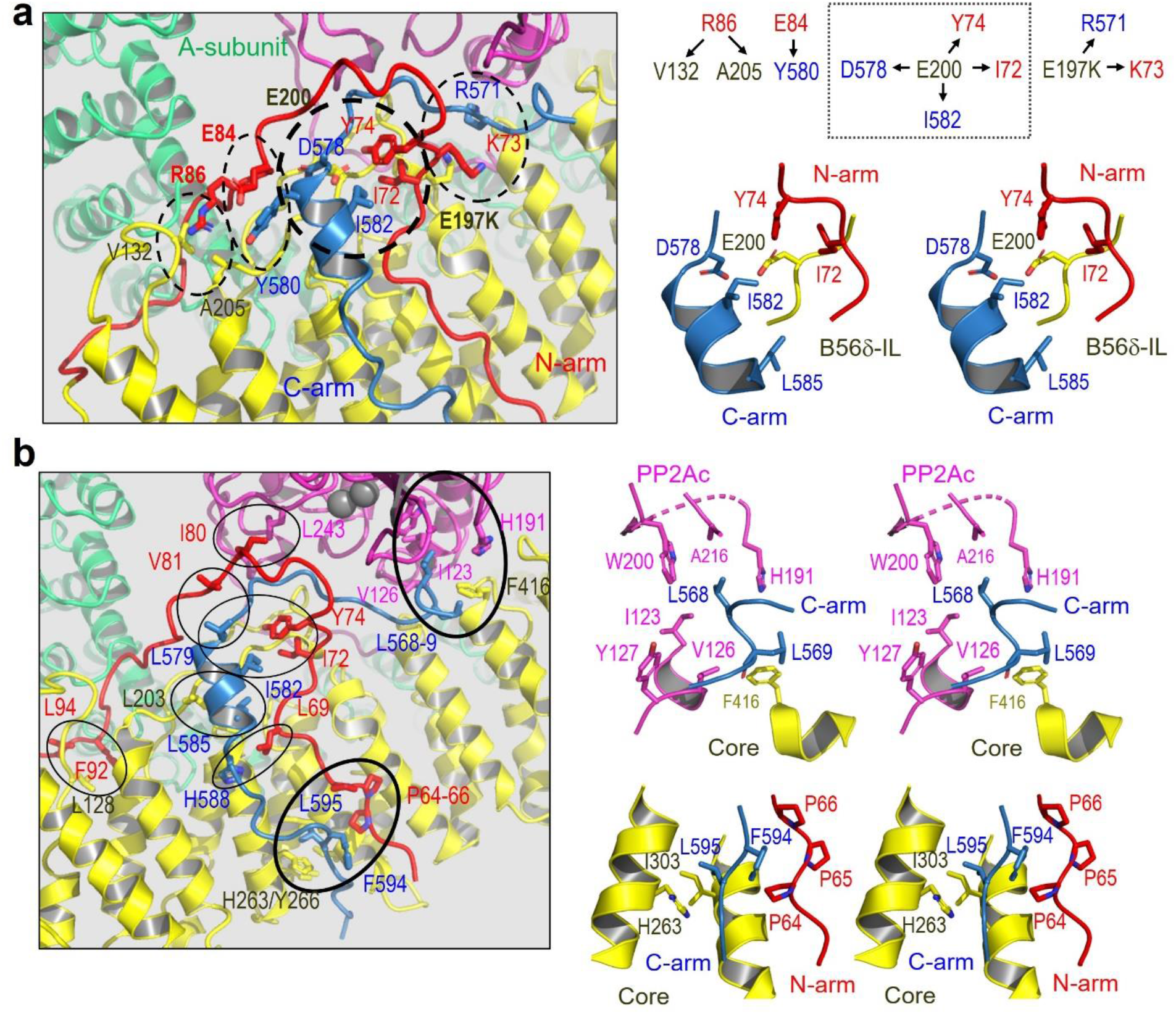
Mapping and closeup views of unfavoarable and hydrophobic contacts along the dynamic interface. (a) Distribution of unfavoarable and repulsive contacts at the dynamic interface, highlighted in dashed cycles (left) and illustrated at the upper right panel. The closeup stereoview of the central repulsive contacts with E200 is shown at the lower right panel. (b) Patches of hydrophobic contacts are in cycles and major hydrophobic interfaces are highlighted in thick cycles (left). The closeup stereoviews for the latter are shown (right). For (a-b), the color scheme is the same as in Fig.1. The structural models are shown in cartoon and key residues at the interfaces in sticks.

Here we determined a high-resolution cryo-EM structure of a PP2A-B56δ holoenzyme bearing the E197K disease mutation at 2.7 Å. The structure resembles a closed form of the holoenzyme. The long N/C-extensions of B56δ make cross-arm interactions against each other and the holoenzyme core, creating an extended dynamic interface that suppresses both the phosphatase active site and the substrate SLiM-binding groove. Nearly all of the previously identified phosphorylation sites, as well as the residues mutated in individuals with intellectual disabilities (ID), are located at or near this interface. We further demonstrated that ID mutations alter the activation phosphorylation rates in response to cAMP-induced activation of PKA. ID mutations also alter the basal level of SLiM-binding and up to a quarter of critical cellular signaling endpoints that affect cell cycle progression and mitotic defects during cell division that could help explain macrocephaly observed in humans. Our studies reveal a coherent allosteric network crucial for phosphorylation-mediated B56δ holoenzyme activation that is altered globally by ID mutations.

## Results

### Overall cryo-EM structures of the PP2A-B56δ holoenzyme

Structure determination of the PP2A-B56δ holoenzyme by single-particle cryo-EM turned out to be highly challenging. We explored the “spotiton” technology^45, 46^ to capture the dynamic states of the holoenzyme on the grids. The spotiton grids “purified’ a closed form of the holoenzyme by dissociating the majority of the holoenzyme particles (Extended Data Fig. 3a). Albeit this form represents a small fraction of total particles, it gave a 4 Å map with uniform density for the long disordered regions at the N/C-termini (Extended Data Fig. 3b). We further explored glutaraldehyde and EDC (ethyl carbodiimide) crosslinks and tested different detergents and EM grids to control ice thickness and particle behavior. Nonetheless, the cryo-EM data for the crosslinked holoenzyme failed to capture any closed forms. Next, we examined the holoenzyme bearing different ID mutations. After an extensive effort, we revealed two major forms for the E197K variant of the holoenzyme: a closed form with a map of 2.59 Å and a loose form with a map of 3.13 Å (Extended Data Fig. 4). The crystal structure of the PP2A-B56γ1 holoenzyme (PDB code: 2NYL)^47, 48^, which represents the common core for the B56 family (Extended Data Fig. 1), fits the map for the B56δ holoenzyme core quite well. Since the N/C-arms are invisible in the loose form, we next focused on the structure of the closed form of the holoenzyme.

**Figure 3.**
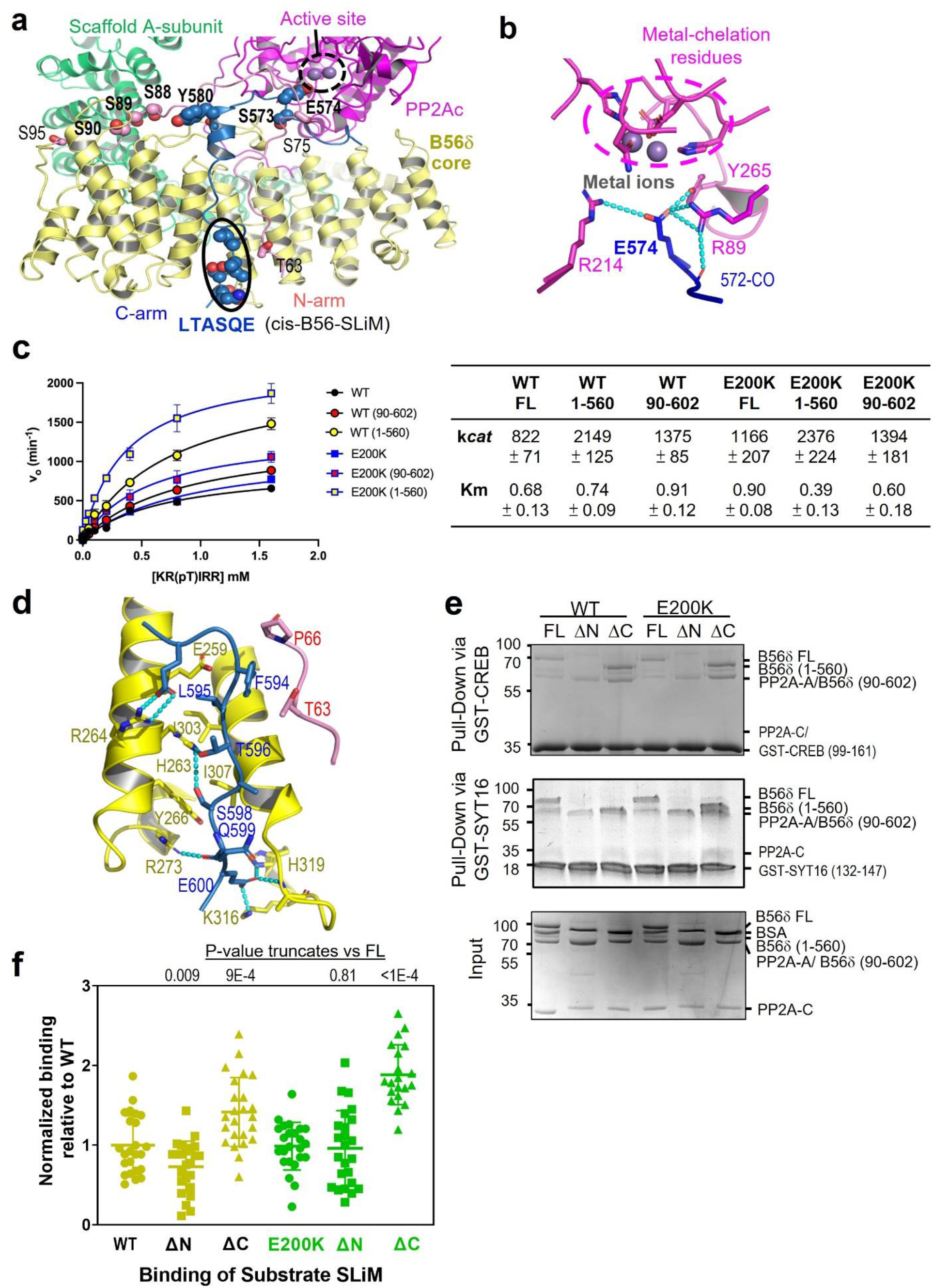
Structural mechanism of dual auto-inhibition and roles of the B56δ N/C-arms. (a) The overall structure of the PP2A-B56δ holoenzyme highlights residues on the N/C-arms essential for activation phosphorylation and suppressing the phosphatase active site and the SLiM-binding groove. The structure is shown in cartoon and colored as in Fig. 1, except that the N/C-arms are colored pink and blue, respectively. Residues with key regulation functions and manganese ions (grey) are shown in spheres. (b) The closeup view of auto-inhibition at the phosphatase active site. The active site metal ions are in spheres. Active site residues and E574 from the C-arm are shown in sticks. (c) Truncations of either N- or C-arm increase the phosphatase activity of both the WT and E200K holoenzymes. (d) The closeup view of auto-inhibition at the B56 SLiM-binding groove, buttressed by extended interactions. (e) Examples of pulldown assays of the WT and E200K PP2A-B56δ holoenzyme full length (FL), truncation of N-arm (ΔN) or C-arm (ΔC) via GST-tagged CREB (99-161) (upper) or GST-SYT16 (132-147) (middle). One fifth of the holoenzyme input is shown (lower). (f) All experimental repeats from (e) are normalized to the WT holoenzyme, and the scatter plots of the normalized results, averages of all repeats, and standard deviation (SD) are shown. The P-values for the full-length versus truncated holoenzymes are calculated using Welch’s t test. For (b) and (d), the structural models are shown in cartoon, and the color scheme is the same as in (a). Residues at the interfaces are shown in sticks. H-bond interactions are shown in cyan dashed lines.

**Figure 4.**
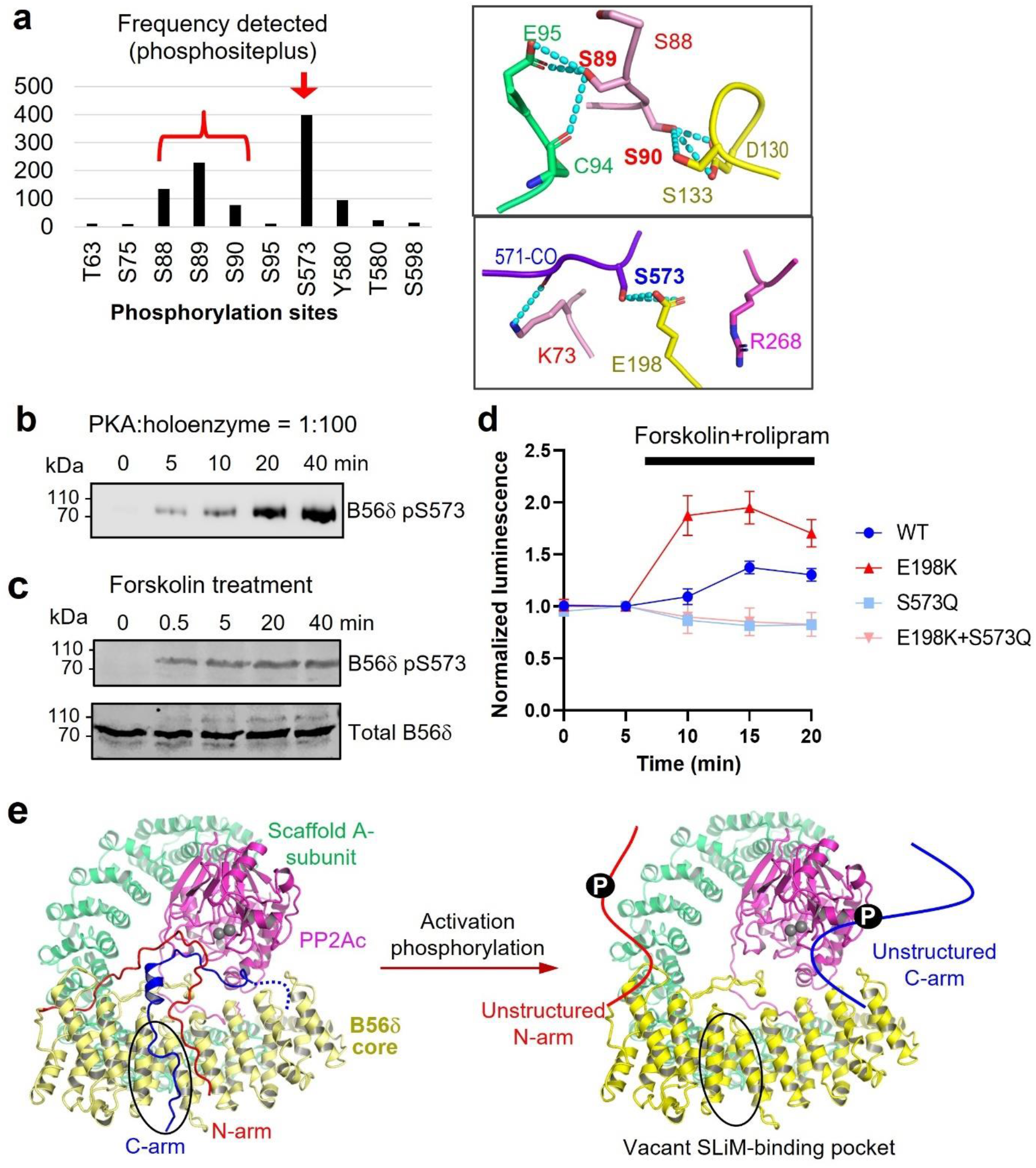
Structural mechanisms of activation phosphorylation of the PP2A-B56δ holoenzyme. (a) The most frequently detected phosphorylation sites on the N/C-arms are highlighted (as summarized from phosphosite.org) (left). The closeup views of these serine residues on the N/C- arms and their interactions at the dynamic regulation interface (right). The color scheme is the same as in Fig. 3a. Phosphorylation at these sites is expected to create repulsive contacts. (b) Time-dependent changes in pS573 of the PP2A-B56δ holoenzyme by PKA *in vitro*. (c) Time-dependent increase of B56δ pS573 upon cellular activation of cAMP/PKA. (d) Time-dependent increase of substrate B56 SLiM binding the PP2A-B56δ holoenzyme (WT and E198K) in mammalian cells upon cellular activation of cAMP/PKA by forskolin and rolipram. The non-phosphorylatable mutation of B56δ, S573Q, abolishes this response. (e) Structural illustration of phosphorylation-induced loosening of the N/C-arms for holoenzyme activation. Phosphorylation of N/C-arms disrupts their interactions with the holoenzyme core, resulting in dual activation of the phosphatase active site and the B56 SLiM-binding groove. The color scheme is the same as in Fig. 1.

The building of the N/C-arms in the closed form was guided by XL-MS (crosslink mass spectrometry). Using zero-length EDC crosslink followed by multiple protease digestions and MS runs, two crosslinked residue pairs in the N/C-arms gave detection frequency comparable to or better than those in the holoenzyme core (Extended Data Fig. 5). The structure of the closed form was refined at 2.7 Å (Extended data Table 1 and Extended Data Fig. 4 and 6). It reveals extensive cross-arm interactions between the disordered regions of the B56δ termini; they interact with the holoenzyme core along a long path that spans over 190 Å (Fig. 1). While the long, linear interface involving two remote disordered arms is intrinsically dynamic in nature, it also harbors a significant number of unfavorable contacts (Fig. 1b), underlying an unprecedented super-long dynamic interface in this highly regulated holoenzyme.

**Figure 5.**
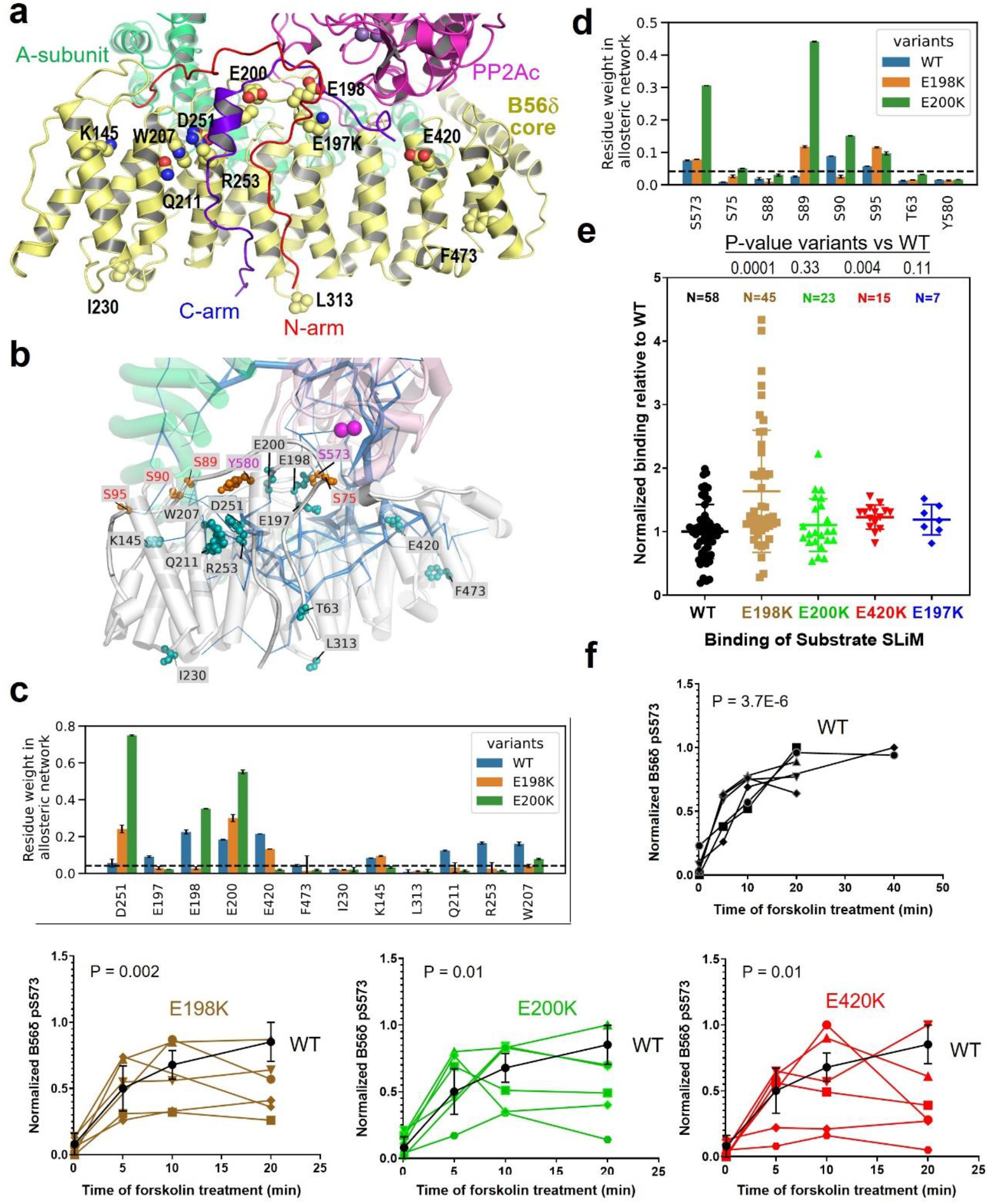
The allosteric network of the PP2A-B56δ holoenzyme, perturbation by B56δ ID mutations, and effects on holoenzyme functions. (a) The overall structure of the B56δ holoenzyme highlights the B56δ ID residues predominantly located at the dynamic regulation interface. (b) Illustration of the allosteric network of the PP2A-B56δ holoenzyme and its relationship to ID residues (green ball-and-stick) and activation phosphorylation sites on N/C- arms (orange ball-and-stick). The allosteric weights are shown as the thickness of blue wires. They are estimated from REST2 simulations of the WT holoenzyme, modified from the cryo-EM structure of the E197K holoenzyme in the closed form. (c-d) The global perturbation of residue weights of ID residues and activation phosphorylation sites on the allosteric network by E198K and E200K. The results in (b-d) are the average of 20 REST2 trajectories. (e) Pulldown of WT and mutant holoenzymes by GST-tagged CREB (99-161) assessed the effects of ID mutations on substrate SLiM binding. The data is normalized with the binding intensity of WT. The number of repeats, their scatter plots, averages, and SD are shown. The P values for comparison of WT and disease variants are calculated using Welch’s t test. (f) Time-dependent phosphorylation of S573 in HEK293 cells expressing WT and mutant B56δ upon cellular cAMP/PKA stimulation. The experiments were repeated six times. Means ± SD was calculated for WT for comparison to disease variants. The P-values for time-dependent changes were calculated using Jonckheere- Terpstra test.

**Figure 6.**
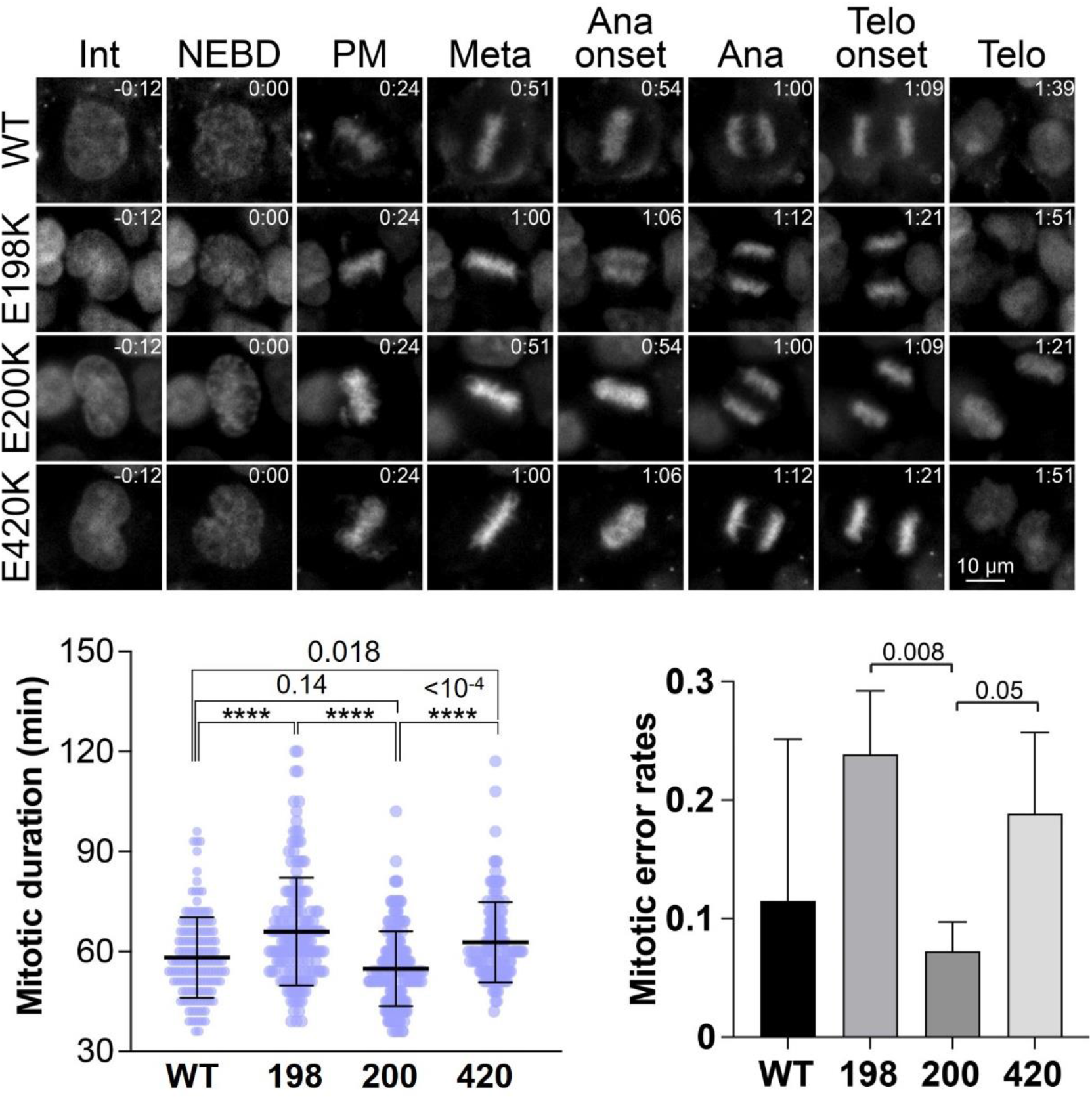
Effects of B56δ ID mutations on mitotic progression and mitotic errors. Representative live cell images of WT and B56δ ID mutant cells (E198K, E200K, and E420K) in HEK293 cells (top). Mitotic durations from NEBD to anaphase onset (bottom left) and the frequency of mitotic errors (bottom right) in the above cells are plotted. Means ± SD was calculated and shown. The P values for comparison of WT and disease variants are calculated using One-Way Anova (bottom, left) and two-tailed t-test (bottom, right). P-values <0.0001 are indicated by “****”. N = 148, 162, 183 and 132 for WT, E198K, E200K and E420K cells.

### The super-dynamic interface in the PP2A-B56δ holoenzyme

The above long-distance interface in the PP2A-B56δ holoenzyme centers at the N/C-arm crossover and makes close contacts with the PP2A catalytic subunit (PP2Ac) and the internal loop of B56 (B56-IL) (I) (Fig. 1b and Extended Data Fig. 7a). The N/C-arms are juxtaposed against each other toward the termini and make close contacts with the B56δ core (II) (Fig. 1b and Extended Data Fig. 7b). Diverging from the crossover and perpendicular to the juxtaposed lower N/C-arms, the upper C-arm makes rich hydrophobic contacts with PP2Ac and miscellaneous contacts with the B56δ core and B56-IL (III) (Fig. 1b and Extended Data Fig. 7c); at the other side, the upper N-arm passes through the cleft between the A-subunit and the B56δ core and makes different modes of contacts with both (IV) (Fig. 1b and Extended Data Fig. 7d).

Intriguingly, the N/C-arm crossover is most rich in repulsive contacts (Fig. 1b). Most prominently, E200 in B56-IL makes repulsive contacts to D578 in the C-arm and unfavorable contacts to several hydrophobic residues in the N/C-arms, I72, Y74, and I582 (Fig. 2a). In addition, E84 in the N-arm makes unfavorable contacts with Y580, a hydrophobic residue in the C-arm (Fig. 2a). These repulsive contacts, together with the energetically unfavorable bending of the N/C-arms, make the crossover the “hottest” hubs along the extended dynamic interface.

Other unfavorable contacts involve a nearby residue R86 in the upper N-arm (Fig. 2a). The E197K ID mutation creates two repulsive contacts to R571 and K73 in the C-arm and N-arm, respectively (Fig. 2a). Such changes might reduce the number of conformational states of the holoenzyme, and thus allowed us to determine the structure of the closed form of the holoenzyme using single-particle cryo-EM.

We further mapped the hydrophobic contacts along the dynamic interface (Fig. 2b). While many small patches of hydrophobic contacts intertwine with the above unfavorable contacts, the dominant rich hydrophobic contacts are made by the two visible ends of the C-arm. L568 and L569 in the upper C-arm are nestled in a hydrophobic pocket formed by six PP2Ac residues and make close contacts with F416 from the B56δ core (Fig. 2b, upper right). Near the C-arm’s terminus, L595 interacts with I303 and H263 at the B56 SLiM-binding pocket, which is buttressed by contacts between F594 to P64/65/67 in the N-arm (Fig. 2b, lower right). The distinct maps of energetically repulsive and favorable hydrophobic contacts likely dictate the complex regulation of the holoenzyme functions.

### Dual autoinhibition of the holoenzyme and roles of the N/C-arms

The PP2A-B56 holoenzymes target specific substrates via the SLiM-binding pocket and the phosphatase active site that are around 30 Å apart (Extended Data Fig. 8). Phospho-substrates containing B56 SLiMs bind to the SLiM -binding pocket, which in turn places the nearby substrate phosphorylation sites close to the PP2A active site. The closed form of the holoenzyme establishes an elegant mechanism of dual autoinhibition, in which E574 and a cis-B56 SLiM (L595TASQE600) in the C-arm make close contacts to the PP2A active site and the B56 SLiM- binding groove (Fig. 3a). E574 in the C-arm mimics the substrate phosphate and makes extensive H-bond and salt bridge interactions to basic residues at the PP2A active site (Fig. 3b). We hypothesize that autoinhibition of the holoenzyme is established by both N/C-arms. Consistent with this notion, deletion of the C-arm (1-560 or ΔC) in either the WT holoenzyme or the E200K disease variant drastically increases the phosphatase activity, and the truncation of the N-arm (90-602 or ΔN) also increases the phosphatase activity of both holoenzymes (Fig. 3c). While the binding of the cis-B56 SLiM to the SLiM-binding groove mimics the substrate SLiMs from substrates^19, 26^, it is also buttressed by the hydrophobic contacts between the upstream F594 with the N-arm (Fig. 3d). Since the holoenzyme is rapidly denatured during isothermal titration calorimetry (ITC), most likely due to the super-dynamic nature of the N/C-arms, we assessed the binding of the GST-tagged substrate SLiMs from CREB^33^ and SYT16^17^ to the full-length and truncated holoenzymes using pulldown assay (Fig. 3e). Consistent with the dynamic nature of the N/C-arms, we noticed a significant fluctuation in the experimental results. Therefore, we repeated the experiments 25 times using three batches of materials to learn about the dynamic range of the substrate SLiM-binding (Fig. 3f). The calculated P-values showed significant differences between truncated and full-length holoenzymes. Intriguingly, while the truncation of the N-arm increases the phosphatase activity (Fig. 3c), it significantly decreases the binding of substrate SLiMs (Fig. 3e-f). Our data suggested that the N/C arms confer coherent suppression at the phosphatase active site but opposing roles at the substrate SLiM-binding groove. It is likely that the kink on the C-arm stabilized by the N-arm is required for the suppression of the active site but causes structural tension toward the SLiM-binding groove (Fig. 3a). In the absence of the N-arm, the C-arm binds the SLiM-binding groove tighter with a relaxed kink. Its multipartite contacts with the holoenzyme core, particularly hydrophobic contacts, are largely intact (Fig. 2b).

### Activation phosphorylation

In addition to the structural modalities for dual autoinhibition, the N/C-arms are rich in phosphorylation sites (Fig. 3a). Among these sites, S88, S89, S90, and S573 are the most frequently phosphorylated sites according to the PhosphositePlus database (phosphosite.org). S89, S90, and S573 make extensive H-bond interactions with acidic residues in the holoenzyme core, namely E95 from the scaffold A-subunit, D130 in the B56δ core, and E198 in the B56-IL (Fig. 4a). Phosphorylation at all these sites is expected to create repulsive contacts that would disrupt the closed form and stimulate the loosening of the holoenzyme, leading to its activation. Consistently, phosphorylation of mouse B56δ at S566 (corresponding to S573 of human B56δ) was found to be associated with the activation of the holoenzyme^40^. Using an antibody that we developed to specifically recognize B56δ phosphorylation at S573 (Extended Data Fig. 9), we demonstrated a time-dependent increase of pS573 *in vitro* upon co-incubation of PKA and the holoenzyme (Fig. 4b) and in HEK293 cells upon forskolin-induced cAMP/PKA activation (Fig. 4c).

To detect the conformational changes associated with activation phosphorylation, we developed a split NanoBiT B56δ holoenzyme sensor, in which the SmBiT peptide fragment was inserted in the holoenzyme core immediately downstream of the N-arm and the LgBiT fragment was fused to the C-terminus of the C-arm (Extended Data Fig. 10). The two NanoBiT fragments are spatially separated in the closed form. If the N/C-arms loosen upon activation phosphorylation by PKA, the two fragments can interact and form an active NanoBiT enzyme (Extended Data Fig. 10). By expressing the WT and E198K mutant holoenzymes bearing this holoenzyme conformation sensor in COS-1 cells, we showed that both holoenzymes produced increased NanoBiT luciferase activity in response to forskolin and rolipram, the combination of an adenine cyclase activator and a PDE4 inhibitor that increases intracellular cAMP levels (Fig. 4d). The responses of both WT and E198K were abolished by S573Q, corroborating the notion that pS573 is essential for holoenzyme activation.

Both our structural observations and biochemical data support a model of holoenzyme activation upon phosphorylation of the N/C-arms (Fig. 4e). Prior to phosphorylation, the holoenzyme prefers a close conformation with crossover interactions of the N/C-arms to establish dual suppression of both the PP2A active site and the B56 SLiM-binding pocket. Upon phosphorylation of the N/C-arms, the repulsive contacts created at the phosphorylation sites drive the N/C-arms from the close conformation to loosen conformations, allowing access to both the phosphatase active site and the B56 SLiM-binding pocket.

### Effects of ID mutations on the holoenzyme allosteric network, autoinhibition, and activation phosphorylation

We reasoned that the super-dynamic interface in the PP2A-B56δ holoenzyme forms a coherent allosteric network connecting the activation phosphorylation sites and the structural elements for autoinhibition. Intriguingly, nearly all ID residues found in Jordan Syndrome patients are spread throughout this dynamic interface (Fig. 5a). These mutations could allosterically impact the dissociation of the N/C arms to alter the holoenzyme activity. To gain critical insights into this allosteric network, we performed molecular dynamic (MD) simulations of the closed form of the holoenzymes for WT and two highly recurrent variants, E198K and E200K. Our initial ten 100- ns MD simulations were not successful in sampling the dissociations of the N/C arms. We thus adopted an enhanced sampling technique: replica exchange with solute tempering-2 (REST2)^49^, in which we included the terminal 8 residues of the N/C-arms in a “hot region”. Twenty replicas were run spanning the temperature space from 310K to 600K. These REST2 simulations resulted in a remarkable improvement in conformational sampling, which led to notable fluctuations in the tails and exposed the B56 SLiM-binding pocket, although complete opening of the arms was not observed (Extended Data Fig. 11a and b).

The REST2 conformations from the lowest temperature replica were used to compute the allosteric network. The importance of a particular residue in the network was calculated as the number shortest paths that cross the residue and normalized by the maximum. A residue was evaluated as important if its weight in the network, was above a cutoff, defined as 80% of all residue weights in the network (Extended Data Fig. 11c, black dashed line in Fig. 5c and d).

Intriguingly, all ID residues in the B56δ core and phosphorylation sites on the N/C-arms are closely integrated into the allosteric network of the holoenzyme (Fig. 5b). Residues F473, I230, and L313 are located on the periphery, and their weight in the network is below the threshold. Either the E198K or the E200K ID mutation alters the residue weights of other ID residues and phosphorylation sites in the allosteric network (Fig. 5c-d), suggesting that ID mutations globally perturb the coherent dual autoinhibition and phosphorylation-induced holoenzyme activation.

Assessing the effects of ID mutations on the holoenzyme function, including substrate SLiM binding and activation phosphorylation rates, turned out to be very challenging. The reproducibility between experimental repeats is considerably low. Given the super-dynamic nature of the B56δ holoenzyme, we initiated an effort to critically control the purification procedures, aiming to precisely control the holoenzyme behavior. We tested two different strains of Hi5 host cells, varied the salt concentrations between 50 mM and 150 mM, and compared French press and Dounce in cell disruption. These variations did not lead to any improvement in the holoenzyme behavior and data fluctuation. No difference was observed in the basal level of holoenzyme phosphorylation, either. The super-dynamic nature of the holoenzymes was also reflected by the fact that the holoenzymes rapidly denatured and formed aggregates during ITC (isothermal calorimetry) experiments.

Nonetheless, summarizing the results from up to ten batches of WT and mutant holoenzymes from the above effort, we observed a statistically significant increase in substrate SLiM binding by the E198K ID mutation despite the range of data fluctuation (Fig. 5e). E420K also shows a substantial effect on the basal substrate SLiM-binding; in contrast, the E200K and E197K variants gave mild effects (Fig. 5e). Next, we assessed time-dependent changes of pS573 upon forskolin-induced cAMP/PKA activation in WT and CRISPR-edited HEK 293 cells bearing homozygotic E198K, heterozygotic E200K, or homozygotic E420K^50^. While the pS573 level of the WT holoenzyme is increased at a stable rate, the phosphorylation rate of the disease variants fluctuates in a large range among all experimental repeats (Fig.5f). Our data support a notion that the tight coupling of the dynamic interface, the holoenzyme allosteric network, the regulatory structural elements, and ID mutations render the disease mutations to differentially modulate the basal activity and the activation phosphorylation rates of the holoenzyme. These highly intertwined structural and functional connections are expected to be highly sensitive to experimental conditions and cellular signaling contexts.

### Effects of B56δ ID mutations on mitosis

B56 SLiMs are predicted in broad signaling proteins involved in different stages of the cell cycle^17^. Building on previous observations of the role of B56δ at the G2/M checkpoint^42^ and mitotic exit^43^, we examined whether the ID mutations E198K, E200K, or E420K (introduced by CRISPR editing) affect the mitotic duration and fidelity of HEK293 cells. We used high temporal resolution live-cell imaging to monitor the mitotic progression in HEK293 cells with either WT or mutant B56δ (Fig. 6). The stages of mitosis were determined using a far-red SiR-DNA dye. The mitotic duration (from nuclear envelope breakdown (NEBD) to anaphase onset) was 58 ± 12 min (N = 148) in the WT cells, but significantly extended in E198K (66 ± 16 min, N = 162) and E420K (63 ± 12 min, N = 132) mutant cells. Intriguingly, E200K mutant cells exhibited similar duration to WT (55 ± 11 min, N = 183) and significantly shorter than E198K and E420 mutant cells. Consistent with this observation, the E200K cells displayed a significantly lower frequency of mitotic errors compared to E198K and E420K cells, the two most severe mutations of the disease variants.

## Discussion

Our cryo-EM structures of the PP2A-B56δ holoenzyme, particularly in its closed form, reveal distinct mechanisms by which it can exist in a dual latent state and become activated in response to phosphorylation. The extended dynamic regulation interface of the holoenzyme provides a coherent framework for understanding the holoenzyme’s dual latency and activation, as well as the shared mechanisms underlying the diverse disease mutations associated with neurological disorders. The complexity of the holoenzyme regulation resides in the unprecedented length of the regulation interface primarily formed by the intrinsically disordered regions rich in regulatory elements. These elements include multiple phosphorylation sites on both N/C-arms, a cis-B56 SLiM and E574 on the C-arm, which establish dual autoinhibition in the closed form, and other signaling sequences on the C-arm, such as nuclear localization signal (NLS) and SH3-binding motif. In addition, the regulation interface is featured by multiple repulsive contacts that contribute to its semi-stable and dynamic nature, which is responsive to phosphorylation and disease mutations. Furthermore, REST2 MD simulation reveals an allosteric network closely coupled to the regulation interface, reflecting the structural intricacy and coherency underlying complex holoenzyme regulation.

The mutation spectrum in patients also corroborates our above notions on the regulation interface in the B56δ holoenzyme. The latter, in turn, provides critical insights into the underlying mechanisms of ID-associated disease mutations. Albeit the mutated residues of diverse variants are widely separated spatially, the super-long regulation interface harbors nearly all of them like an extended umbrella. The coupling of ID mutations to the regulation interface and the allosteric network explains their shared symptoms, including intellectual disability, hypotonia, and autism^8,9, 12, 13^. ID mutations at the holoenzyme regulation interface might perturb the holoenzyme regulation and function in many scenarios, potentially leading to dominant negative effects. For example, ID mutations might perturb the dual latency individually or simultaneously and alter the activation phosphorylation rates of the holoenzyme. Furthermore, the loss of coherent dual latency and perturbed holoenzyme activation might also lead to untimed exposure of NLS and binding sites for SH3 and 14-3-3. Consistent with the latter notion, a family with B56δ missing the C-arm showed incomplete penetrance of neurological disorders compared to other missense variants that are *de novo* and completely penetrant^51^. The effects of B56δ mutations are also supported by studies using CRISPR-edited HEK293 cells bearing heterozygotic or homozygotic

E420K^50^. This mutation in either single or both alleles perturbs the same diversity of signaling nodes.

The involvement of the B56 SLiM and its binding groove at the regulation interface likely contribute to the bulk of the broad range of clinical symptoms of Jordan Syndrome. The human proteome has ∼1,500 B56 SLiM-containing proteins involved in broad cellular signaling and processes^17^. Perturbation of the dual latency and phosphorylation-induced holoenzyme activation is expected to affect many of these B56 SLiM-containing signaling proteins. Consistently, CRISPR-edited HEK293 cells bearing B56δ E420K altered 6% of phosphopeptides in the proteome by at least 2-fold^50^. We showed that among ∼1,000 genes in SFARI Gene database with genes associated with autism and neurodevelopmental disorders (gene.sfari.org), >10% possess B56-targeting SLiMs (data not shown), underlying overlapping molecular processes perturbed by Jordan Syndrome and SFARI genes. In addition to the effects of ID mutations on mitosis (Fig. 6), the E420K mutation was shown to affect mTOR/AKT signaling among other signaling processes^50^. Coupling to PP2A holoenzyme SLiMs is an emerging theme of signaling and functional complexity. For example, the multiple PP2A holoenzyme SLiMs in the PP2A methylesterase (PME-1) creates versatile PME-1 activities toward PP2A holoenzymes, diverse signaling pathways, and cellular processes^31^.

In addition to the shared mechanisms across variants, our study begins to illuminate the molecular basis of the genotype-phenotype correlation of clinical severity. Clinical studies showed that E198K is the most severe form of the disease and that E420K is more severe than the E200K and E197K variants^8, 9, 12^. We showed that the E198K holoenzyme has a much higher basal level of substrate SLiM-binding than the WT holoenzyme and other disease variants (Fig. 5e). Furthermore, E198K and E420K cause longer mitotic duration than WT and E200K and gave higher mitotic error rates than the E200K variant (Fig. 6). In a companion paper focusing on the changes of the holoenzyme allosteric paths by E198K and E200K, we showed that the E198K holoenzyme has an allosteric pathway resembling the holoenzyme with activation phosphorylation, underlying a higher basal activity and loss of latency of this variant^52^. In contrast, the allosteric pathway is minimally affected by E200K. Many questions remain to be answered. Our study laid the critical foundation for understanding the function and regulation of the B56δ holoenzyme and an in-depth understanding of the underlying mechanisms of Jordan Syndrome mutations.

It is important to stress the super-dynamic nature of the regulation interface formed by the disordered N/C-arms and the holoenzyme core. Our challenges to capture the closed form of the WT holoenzyme by single-particle cryo-EM suggests the presence of many conformation states that might be required for numerous nuances of cellular signaling under normal physiological conditions. Two distinct structural features might contribute to the super-dynamic nature: the interactions of two extended intrinsically disordered regions from the widely separated N/C-arms and the multiple repulsive contacts along the interface. Enriching broad regulation elements in the disordered regions forms one more basis for the complexity and intricacy of this regulation interface. The structural and functional features outlined here might not be unique to the B56δ holoenzyme but are a common theme for modulating complex biological processes and signaling that remain to be broadly studied.

## Author Contributions

CGW performed cryo-EM studies, assisted and guided by HW, BC, WKC, and YX. CGW and YX determined the structures. CGW, VKB, and PSP performed biochemical studies, assisted by RS. KK performed REST2 simulation, guided by XH and YX. BEW generated anti-pS573 antibody. MS, AM generated CRISPR cell lines, guided by RH. YCC, CGW, and RS performed cell biology, guided by YX and AS. RAM developed and tested the holoenzyme sensor, guided by SS. YX performed data analysis and structural analysis, developed the figure panels, and wrote the manuscript, assisted by CGW.

## Supporting information

manuscript

## Acknowledgement

We thank Drs Derek Taylor and Wei Huang from Case Western Reserve University for discussions on cryo-EM grid freezing, Drs Janette Myers from PNCC (Pacific Northwest Cryo-EM Center) for assistance and discussions on cryo-EM data collection, and Kenneth Satyshur and Ivan Andrewjeski from School of Medicine for IT support. Cryo-EM data collection was supported by NIH grant U24GM129547, performed at PNCC at OHSU, and accessed through EMSL (grid.436923.9), a DOE Office of Science User Facility sponsored by the Office of Biological and Environmental Research. This study is supported by NIGMS R01 GM137090-01 (YX), Jordan’s Guardian Angels Foundation, and Jordan’s Syndrome research consortium fund from the State of California A19-3376-5007 (YX, sub-award from 2021 SB 129 #44, 2018 SB 840, PD/PI: Nolta, Jan), NIGMS R35 GM147525 (AS), and the Hirschfelder Professorship Fund (XH).

## Materials and Methods

### Protein preparation

All protein constructs were generated by standard PCR molecular cloning strategy. The human His6-tagged scaffold A-subunit (α isoform), His8-tagged PP2Ac (α isoform), and GST-tagged B56δ were overexpressed in insect cells using the lab-modified Bac-to-Bac baculovirus expression system^53, 54^. Briefly, Hi-5 cells grown to a density of 1.5 × 10^6^ cell/ml were co- infected with PP2A His-scaffold A subunit, His-PP2Ac, and GST-B56δ baculovirus for 48h at 27⁰C. Cells were lysed by Dounce homogenization in the lysis buffer containing 25mM Tris-HCl (pH 8.0), 150mM NaCl, 50 μM MnCl2, 2mM DTT, and protease inhibitors (10μM leupeptin, 0.5 μM Aprotinin and 1mM PMSF). The insoluble proteins were removed by centrifugation, and the soluble fraction of cell lysates was gravity-loaded to GS4B (Glutathione Sepharose 4B) resin (Cytiva) column three times followed by two washes using 5 column volumes (CV) of lysis buffer. The proteins left on the resins were digested by the TEV protease, and the flow-through was further fractionated by anion exchange chromatography (Source 15Q column, Cytiva) and gel filtration chromatography (Superdex 200 column, Cytiva). The point mutations for the B56δ disease variants and/or truncations of the B56δ N/C-arms were introduced by site-directed mutagenesis, and the mutant and/or truncated holoenzymes were expressed and purified similarly to the WT holoenzyme. GST-tagged SYT16 (132-147) and CREB (99-161) were overexpressed in *E. coli* DH5α and purified over GS4B resin and ion exchange chromatography.

### GST-mediated pulldown assay

To test the interaction between WT and mutant PP2A-B56δ holoenzyme with substrate B56 SLiM peptides, 12 μg of GST-tagged SYT16 (132-147) or GST-tagged CREB (99-161) was immobilized on 5 μl of GS4B resin. The unbound protein was washed out by assay buffer containing 25 mM Tris (pH 8.0), 150 mM NaCl, 3 mM DTT, and 1x protease inhibitor cocktail (P8340, Sigma). 10 μM of PP2A-B56δ holoenzymes were then added to the immobilized GST-tagged protein in a final volume of 50 μl assay buffer supplemented with 1mg ml^-1^ of BSA. After 5 min of incubation, the unbound proteins were removed, and the resins were washed three times with the assay buffer supplemented with 0.1% of Triton X-100. The proteins that remained on the resin were examined by SDS-PAGE and visualized by Coomassie blue staining.

### Cross-linking mass spectrometry (XL-MS) of the PP2A-B56δ holoenzyme

The intra-molecular interactions in the PP2A-B56δ holoenzyme were probed by EDC zero-length chemical crosslinker, followed by mass spectrometry analysis. In brief, 1.5 μM PP2A- B56δ holoenzyme in 25 mM MES (pH6.0) and 150 mM NaCl was incubated with 60 mM EDC at RT for 75 min, followed by quenching the reaction with 100 mM Tris. The crosslinked samples were desalted by desalting columns (Zepa spin desalting column, Thermo Fisher Scientific) and analyzed by SDS-PAGE. The bands representing holoenzymes with inter-subunit crosslinks were excised from SDS-PAGE for reduction and in-gel digestion. The excised bands were reduced in 25mM DTT and 25mM NH4HCO3 at 57⁰C for 30 min and alkylated with 55mM IAA in 25 mM NH4HCO3 in darkness for 30 min at RT, followed by digestion in 20 μl of 10ng ul^-1^ Trypsin (Promega) in 25 mM NH4HCO3 (pH 8.0-8.5) and 0.01% ProteaseMAX w/v (Promega) for 16 hr at 37⁰C. To improve peptide detection and sequence coverage, secondary digestion was performed with 20ng μl^-1^ GluC or chymotrypsin in 25 mM NH4HCO3 for 8 hr at 37⁰C and quenched by 0.05% TFA, followed by desalting with C18 cartridges. The resulting samples were then separated and analyzed by LC-MS/MS using Lumos mass spectrometer. The identification of peptides for all three subunits was conducted by MeroX software^55^. Multiple XL-MS experiments were performed, and four with 90% sequence coverage were used to identify crosslinked residue pairs. The detection frequency for reliable residue pairs was evaluated using the identical pairs in the crystal structure of the PP2A-B56γ1 holoenzyme (PDB: 2NPP).

### Cryo-EM sample preparation and data acquisition

The cryo-EM grids for the WT PP2A-B56δ holoenzyme were prepared at the New York Structural Biology Center using a prototype of the commercial Chameleon system (SPT Labtech) based on the Spotiton technology^45, 46^. 50 pl of the WT holoenzyme at a concentration of 1.6 mg ml^-1^ was applied to the homemade self-wicking nanowire grids^56^ using a piezo-electric dispenser, followed by plunge-frozen in liquid ethane. The robot chamber was operated at room temperature without strictly controlled but moderate humidity. The cryo-EM data for the WT holoenzyme was collected using Titan Krios (ThermoFisher) electron microscope operated at 300 kV equipped with Gatan K2 summit cameras. A total of 1790 movies were automatically acquired by Leginon^57^ with a defocus range of -1.2 to -2μm at a nominal magnification of 105,000×, corresponding to a pixel size of 1.096 Å/pixel. Each stack dose-fractioned over 50 frames was recorded with a total electron dose of 66.84 eÅ^−2^.

For the E197K PP2A-B56δ holoenzyme, an aliquot of 3 μl of purified holoenzyme at 0.4 mg ml^-1^ was applied onto a glow-discharged holey gold grids (UltraAuFoil R1.2/1.3), blotted for 4 s with a blot force of -5 and plunge frozen in liquid ethane using a FEI Vitrobot Mark IV (Thermo Fisher Scientific) at 4°C and 100% humidity. Cryo-EM data were collected using a Titan Krios operating at 300 kV with a Gatan K3 detector and GIF Quantum energy filter. A total of 9158 movies were collected using SerialEM, with a slit width of 20 eV on the energy filter and a defocus range from -0.7 to -2.2 µm in super-resolution counting mode at a nominal magnification of 81,000×, corresponding to a pixel size of 1.068 Å/pixel. Each stack dose- fractioned 69 frames was recorded for a total exposure time of 3.2 s and electron dose of 49 eÅ^−2^.

### Cryo-EM data processing

All data were processed with similar strategies and procedures using cryoSPARC 3.0^58^. Movies were motion-corrected using Patch Motion Correction, followed by the contrast transfer function (CTF) estimation using Patch CTF Estimation. Images with bad CTF estimations worse than 4Å were discarded. Particles were first picked from thirty good images using Blob Picker and extracted with a box size of 280Å. Good 2D class averages showing projections in different orientations were selected as templates for automatic picking for the entire dataset. Reiterate 2D classification was used to remove bad particles. The remaining good particles were used for ab initio reconstruction, followed by heterogeneous and homogenous refinements, with particles from bad classes removed in each reiterate procedure. Different conformations of the PP2A- B56δ holoenzyme were best separated by 3D variability analysis (3DVA)^59^. Particles from conformationally homogeneous classes were then subjected to homogenous refinement, followed by local and global CTF refinement. Local refinement with a mask around the close conformation of the B56δ subunit was performed to obtain a better resolution for this subunit.

These efforts gave maps with a resolution of 3.13 Å for the loose form and 2.59 Å for the closed form of the E197K PP2A-B56δ holoenzyme. The resolution was estimated by applying a soft mask around the protein complex and using the gold-standard Fourier shell correlation (FSC) = 0.143 criterion.

### Model building and refinement

The initial model of the PP2A-B56δ complex was built on the structure of PP2A-B56γ1 holoenzyme (PDB ID: 2NPP) and manually docked into the cryo-EM maps in Chimera^60^. Modifications to the holoenzyme core and model building of the N/C-arms were performed in COOT^61^. The structural model was refined using the phenix.real_space_refine program in PHENIX^62^ with secondary structure and geometry restraints. The final models were analyzed using MolProbity^63^.

### Phosphatase assay

The enzyme kinetics of the purified PP2A-B56δ holoenzymes were determined using the PiColorLock phosphatase assay (Abcam, Ab270004), measuring the release of inorganic phosphate from the dephosphorylation of substrates. A phosphorylated hexapeptide (KRpTIRR) was used as the substrate for the assay. 50ul of 30nM indicated PP2A-B56δ holoenzymes in the assay buffer, containing 25mM Tris pH8.0, 150mM NaCl, and 50uM MnCl2 supplemented with 0.05 mg ml-1 BSA, were added to the 10μl substrate (6 times of the final concentration prepared in the assay buffer) in a 96-well clear plate to start the reactions. After 3 min, the reactions were quenched by 15 μl of quench buffer provided in the kit and allowed the color to develop for 15 min. The absorbance at 635nm of the reactions was then read by the SpectraMax Plus 384 Microplate Reader (Molecular Devices) using the end-point mode. Initial velocity (V0) determined at varying concentrations of substrate was calculated and fit to the Michaelis–Menten equation (eq. 1) to determine the steady-state kinetics of the PP2A-B56 holoenzymes.

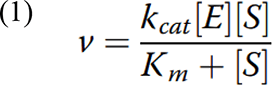

In eq. 1, kcat is the rate constant, [E] and [S] are the enzyme and substrate concentration, and Km is the Michaelis-Menten constant reflecting the binding affinity between the peptide substrate and the enzyme.

### *In vitro* holoenzyme phosphorylation by protein kinase A (PKA)

100 μl PP2A-B56δ holoenzyme was prepared to a final concentration at 0.1mg ml^-1^ in assay buffer containing 50 mM Tris-HCl (pH7.5), 10 mM MgCl2, 200 uM ATP, 0.1 mM EDTA, 2 mM DDT, and 0.01% Brij35. Phosphorylation was initiated by adding 1 μl PKA at 0.1mg ml^-1^ to the PP2A-B56δ holoenzymes (PKA: PP2A-B56δ = 1: 100 (w/w)) and incubating the mixture at 37°C. 10 μl PP2A-B56 holoenzyme was taken out at the indicated time from the reaction, mixed with 3 μl of 4x SDS dye (40% Glycerol, 8% SDS, 0.25M Tris-HCL, pH 6.8,0.4M DTT,0.04% bromophenol blue) and immediately heated at 95°C for 3 min to quench the reaction. 0.05 μg of PP2A-B56 holoenzyme from each time point were examined by western blot. To detect the phosphorylation level at S573 of B56δ, we utilized a newly developed antibody that recognizes pS573 (see supplemental information). The total B56δ was detected by a commercial antibody (Invitrogen, MA-26636).

### Mammalian cell culture and the time course of B56δ phosphorylation upon cellular cAMP activation

Human embryonic kidney cells (HEK293T) cells were cultured in Dulbecco’s modified Eagle’s medium (Gibco, Thermo Fisher Scientific, Waltham, MA, USA) with 10% fetal bovine serum (Hyclone, GE Healthcare, Boston, MA, USA), 100 U ml^-1^ penicillin, and 100 μg ml^-1^ streptomycin in a humidified atmosphere at 37 °C with 5% CO2.

Following transit transfection of the Flag-tagged B56δ (WT, E198K, E200K, and E420K), the HEK293T cells were treated with 10uM forskolin (Fsk; Sigma: 344270-10MG) and 1mM isobutylmethyxanthine (IBMX; Cayman:13347) to stimulate cellular cAMP/PKA. The cells were harvested at the indicated time following stimulation and lysed using a buffer containing 20mM Tris-HCl pH7.0, 150mM NaCl, 0.1% Triton X-100, 10uM H89 (PKA inhibitor), 10ug ml^-1^ DNase, protease inhibitor and phosphatase inhibitor (phoSTOP, Sigma-Millipore). Total cell lysates (20 μg) for each time point were examined by Western blot using antibody that specifically recognizes pS573. The levels of pS573 were normalized to the total B56δ detected by a specific antibody (Invitrogen-Thermal Scientific, MA-26636). At least six experimental repeats were performed to determine the fluctuation range of the activation phosphorylation rates. The P-values for the time-dependent increase of pS573 were calculated using Jonckheere- Terpstra test in MSTAT7.0 (https://oncology.wisc.edu/mstat/).

### REST2 simulation

The cryo-EM structure of the closed form of the E197K PP2A-B56δ holoenzyme was used to generate the initial model for the close forms of WT, E198K, and E200K holoenzymes.

Molecular dynamics (MD) simulations were performed using Gromacs^64^ patched with PLUMED 2.8.0 in the Amber ff14SB forcefield^65^. The protonation state of protein residues was determined by propka 3.4.0. The short-range Coulomb interactions were cut off at 0.9 nm, and long-range electrostatics were computed with PME (particle-mesh Ewald). The Lennard-Jones interactions were cut off at 0.9 nm. The structure was simulated in a dodecahedron box, with the minimum distance to the surface of the protein of 1 nm. The complex was solvated in TIP3P water (approximately 60,000 water molecules), and sodium ions were added until the system was neutral. The energy of the system was minimized with the steepest descent algorithm for 10,000 steps. Then, 1 ns MD simulations were performed at 310K with the positions of all protein atoms except hydrogen restrained. In these simulations, the temperature was controlled by a V-rescale thermostat with a coupling constant of 10 ps, and pressure set to 1 bar controlled by a C-rescale barostat with a coupling constant of 1 ps. Snapshots used for analysis were spaced by 100 ps.

Conventional MD simulations are not successful in sampling the dissociations of the N/C-arms. We thus have adopted an enhanced sampling technique: REST2^49^, with the terminal 8 residues of the N/C-arms placed in a “hot region”. 20 independent replicas were run with the “hot region” temperatures assigned to a geometric progression: 310, 320, 332, 344, 356, 368, 381, 395, 409,

423, 438, 454, 470, 486, 503, 521, 540, 559, 578, and 600K. Exchanges between replicas were attempted every 500 steps and the acceptance probability achieved was ∼20%. REST2 simulations were conducted up to 100 ns, totaling 2µs of simulation time per PP2A variant.

### Allosteric network

The allosteric networks were computed for each variant of the PP2A using an established graph- theoretic approach^66, 67^. To compute a network, each residue in the holoenzyme was represented by two atoms: Cα and the non-hydrogen sidechain atom most distant from the Cα. For residues Ala, Gly and Pro only the Cα atom was used. Then, using all the conformations from the REST2 simulations at T=310K, linear mutual information (LMI) *C*_*ij*_ between all pairs of atoms was computed using g_correlation^68^. Next, the resulting LMI matrix was multiplied with a semi-binary contact map, so that only the neighboring atoms would have a significant contribution to the network. The contact map *K* between each pair of selected atoms is a piecewise smoothing function as used by Botello-Smith *et al*^69^:

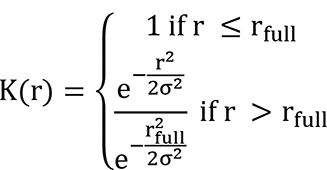

Where *r* is the distance between the selected atoms in an MD snapshot. The cutoff distance for a full contact was selected as *r*_*full*_ = 0.8 nm and *r*_*cut*_ = 1.5 *nm*, so *K*(*r*_*cut*_ = 1.5) = 10^−5^ and 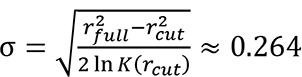. Then, 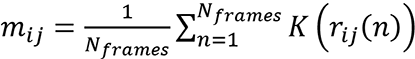 are the elements of the contact map averaged over all frames in the simulation. The LMI matrix with a contact cutoff was used to define the network, where the selected atoms served as nodes, and the edges were equal to the masked LMI. The edge weights in the network were computed as *W*_*ij*_ = −*log*|*L*_*ij*_| where *L*_*ij*_ = *m*_*ij*_ *C*_*ij*_. Paths in this network weighed by the edges explain how strongly the correlated motion of residues in the source affects residues in the sink and were computed with the Dijkstra algorithm using the “networkX” python library (https://networkx.github.io (2020)). Finally, the number of times a residue was crossed by the shortest paths was counted, and contribution from individual atoms in the residue were summed up. This count, normalized by the maximum count in the network, was used as the weight of the residue in the overall allosteric network. Confidence intervals were computed as the standard deviation of residue weight in networks obtained from 10 bootstrapped trajectories. The bootstrapped trajectories of 100 ns were obtained by selecting blocks of 10 ns from the original dataset with replacement.

### Mammalian cell holoenzyme sensor to detect conformation transition upon activation

N-terminal large (LgBit) and C-terminal small (SmBit) fragments derived from NanoLuc (Promega) were fused to the C-terminus of B56δ and inserted downstream of its N-arm at residue 103, respectively. The SmBit sequence is sandwiched by a short linker sequence, “GSG” at both ends to alleviate structural hindrance for complementation with LgBit. The constructs expressing the B56δ holoenzyme conformation sensors were transfected into COS-1 cells. After 24 hours, cells were placed in 20% Nano-Glo® Live Cell Reagent (Promega) mixed with 80% DMEM containing 1% FBS. Cells were equilibrated at 37 °C in 5% CO2 in a Cytation 5 before the luminescence readings were measured every 5 minutes. Following the second reading, cells were treated with 20 μM forskolin (Tocris) and 2 μM rolipram (Tocris) to activate adenylyl cyclase and inhibit cAMP phosphodiesterase 4 (PDE4) simultaneously to activate cellular cAMP/PKA. All values were normalized to the vehicle control, and all wells were normalized to the 5-min time point (just before forskolin/rolipram treatment). The experiments were performed in quadruplicate and repeated three times. Representative results were shown. Error bars represent standard deviation.

### CRISPR prime editing to introduce B56δ ID mutations

A single genomic change of ID mutations was introduced into HEK-293 cells to one or both alleles of *PPP2R5D*. The CRISPR knock-in of the E420K B56δ mutation was produced by cytidine base editing of the coding sequence in exon 12 of *PPP2R5D* as previously described^50^. The heterozygotic and homozygotic E198K knock-ins and the heterozygotic E200K knock-in were produced by prime editing PE3b strategy^70^. Briefly, Cas9 nickase fused to an engineered reverse transcriptase (RT) with improved thermostability and processivity (PE2) was programmed with a prime editing guide RNA (pegRNA) to nick and edit the PAM strand. Once the DNA strap on the edited strand was resolved, a secondary single guide RNA (sgRNA) that matches only the edited strand, not the original allele, guided the Cas9 domain of PE2 to nick the non-edited strand. This strategy, known as PE3b^70^, improved editing efficiency without introducing double-strand breaks and reduced off-target edits. The pCMV-PE2 plasmid expressing PE2 Cas9-RT fusion protein (Addgene plasmid #132775, https://www.addgene.org/132775/) was provided as a generous gift from Dr. David Liu. For each knock-in mutation, a dual RNA expression cassette containing sequences encoding a mutation-specific pegRNA (driven by an hU6 promoter) targeting *PPP2R5D* exon 5, and the secondary mutation-specific sgRNA (driven by a 7SK promoter) was synthesized and cloned into pUC57- kan vector.

Approximately 200,000 wild-type HEK-293 cells were electroporated with pCMV-PE2 and the appropriate RNA expression plasmid using a Neon transfection system for each knock-in mutation. After 48h of recovery, electroporated cells were clonally isolated by single-cell sorting into 96 well plates using a BD FACS Aria II. After clonal expansion, genomic DNA was isolated, and regions containing exon 5 were PCR-amplified. Sanger sequencing was then employed to detect the desired single-base mutations. Before further use, cell lines with the desired mutations were single-cell sorted three times to ensure each cell line represents a homogenous population derived from a single gene-edited cell. Complete genomic sequencing of the parental, E420K, E200K, and E198K variant cell lines was performed to characterize the genomic background, detect any off-target editing, and ensure that repetitive single-cell sorting did not introduce spontaneous somatic mutations. Cell lines with off-target mutations detected in protein-coding regions were discarded. Details for the E198K prime edited cell lines are described elsewhere^71^. The design of pegRNAs and sgRNAs for E200K is described in supplemental materials.

### Live cell imaging and mitotic duration measurements

To investigate whether the B56δ ID mutation, E198K, E200K, or E420K affects the mitotic duration and fidelity, we used high temporal resolution live-cell imaging to monitor the mitotic progression of HEK293T cells with the WT B56δ or CRISPR knock-in ID mutation. Nikon Ti-E inverted microscope equipped with a Photometrics CoolSnap Hq2 CCD camera, spectra-X LED light source (Lumencor), and a Plan Apo 20x objective (NA = 0.45) controlled by Nikon Element software was used for live imaging. Time-lapse imaging collected 5 frame 3D stacks at 2 μm steps along the z-axis at 3 min intervals for 24-48 hours. All cells were grown on a 4- chambered glass bottom dish (#1.5 glass, Cellvis) in FluoroBrite DMEM media (Thermo Fisher) supplemented 10% FBS (Gibco) and 2 mM GlutaMAX (Gibco). All cells were incubated with sirDNA and 10 µM of verapamil (Cytoskeleton, Inc.) for 4 hours before live imaging. Cells were recorded at 37°C with 5% CO2 in a stage-top incubator using the feedback control to maintain the growing media’s temperature (Tokai Hit, STX model). Image analysis was performed using Nikon Element software. Mitotic stages were determined by nuclear staining. The mitotic duration was measured from nuclear envelope breakdown (NEBD) to anaphase onset. Incidences of lagging chromosomes were analyzed. The experiments were independently repeated three times, and P-values between variants were calculated by One-Way Anova and two-tailed t-test. P-values < 0.05 were considered significant.

## Notes

### Competing Interest Statement

The authors have declared no competing interest.

### Summary of Updates

Figure 5 and the corresponding paragraph were updated to improve the clarity Several typos in the previous version were corrected

