## Supplementary material for "Extended regulation interface coupled to the allosteric network and disease mutations in the PP2A-B56δ holoenzyme": manuscript

**Supplemental Materials and Methods**

**Extended Data Figure 1-10 and Table 1**

**References for Supplemental Information**

### Supplemental Materials and Methods

#### Generation of pS573 B56δ antibody

Phospho-specific antibodies recognizing the pS573 epitope in B56δ were developed by adaptation of a previously described protocol[1]. Briefly, peptides encompassing the non-phosphorylated S573 (CLLRRKSELPQ) and phosphorylated S573 (CLLRRKpSELPQ) epitopes were synthesized with a C-terminal Cys residue and conjugated to Imject™ Maleimide-Activated mcKLH (pS573-KLH), Imject™ Maleimide-Activated BSA (S573-BSA and pS573-BSA), and SulfoLink™ Coupling Resin following the manufacturer's recommended protocol (ThermoFisher Scientific). The pS573-KLH conjugate was used for antibody production in alpacas (Turkey Creek Biotechnology, Waverly, TN). Phospho-specific antibodies recognizing the pS573 epitopes were purified from the alpaca antisera by differential affinity chromatography. Briefly, 5 ml of pS573 antisera was passed over the non-phosphorylated peptide column (2 ml). The non-bound material was then applied to the phosphorylated peptide column (2 ml). After washing with 40-50 ml PBS, bound antibodies were eluted with 8.5 ml of 100 mM glycine, pH 2.5, and collected in tubes containing 1.5 ml of 1.5 M Tris-HCl, pH 8. The specificity of the purified pS573 antibodies was then assessed by dot blot analysis using S573-BSA, pS573-BSA, and non-phosphorylated and phosphorylated peptide-BSA conjugates corresponding to other phosphorylation sites in B56δ (Extended data Fig. 9).

#### CRISPR prime editing design for B56δ E200K mutation

The heterozygotic heterozygotic E200K knock-in were produced by prime editing PE3b strategy[2]. Briefly, Cas9 nickase fused to an engineered reverse transcriptase (RT) with improved thermostability and processivity (PE2) was programmed with a prime editing guide RNA (pegRNA) to nick and edit the PAM strand. Once the DNA strap on the edited strand is resolved, a secondary single guide RNA (sgRNA) that matches only the edited strand, not the original allele, will guide the Cas9 domain of PE2 to nick the non-edited strand. In addition to the common sgRNA scaffold sequence, pegRNAs incorporate a target (protospacer) sequence, a primer-binding site (PBS), and a reverse transcriptase template (RTT) encoding the desired mutation. The designs for the E200K knock-in are: pegRNA protospacer = AGGGGCUGAGUUUGACCCAG, PBS = GGUCAAACUCAGC, RTT = GCUUAUCUUCCUCGG. The secondary nicking sgRNAs contain a protospacer that matches the edited strand, not the original allele. The design for E200K sgRNA protospacer = GGGUGGGCUUAUCUUCCUCG.

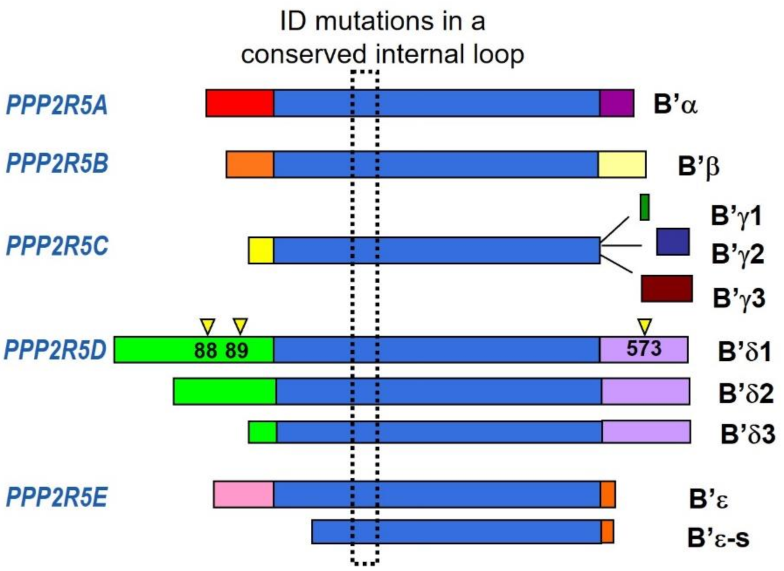


**Extended Data Figure 1. Domain structures of the B56 family of PP2A regulatory subunits show conserved common core (blue) and the unique N- and C-terminal extensions.** B56δ (B56δ1, the longest splicing variant encoded by *PPP2R5D*) has long N- and C-extensions enriched with phosphorylation. B56γ1, the shorted splicing variant encoded by *PPP2R5C*, represents the common core of the B56 family. The structural region (dashed box) consists of a conserved internal loop in the B56 family of PP2A regulatory subunits is a hot spot where many ID mutations are found in Jordan's syndrome. The most frequently detected phosphorylation sites are highlighted with yellow triangles.

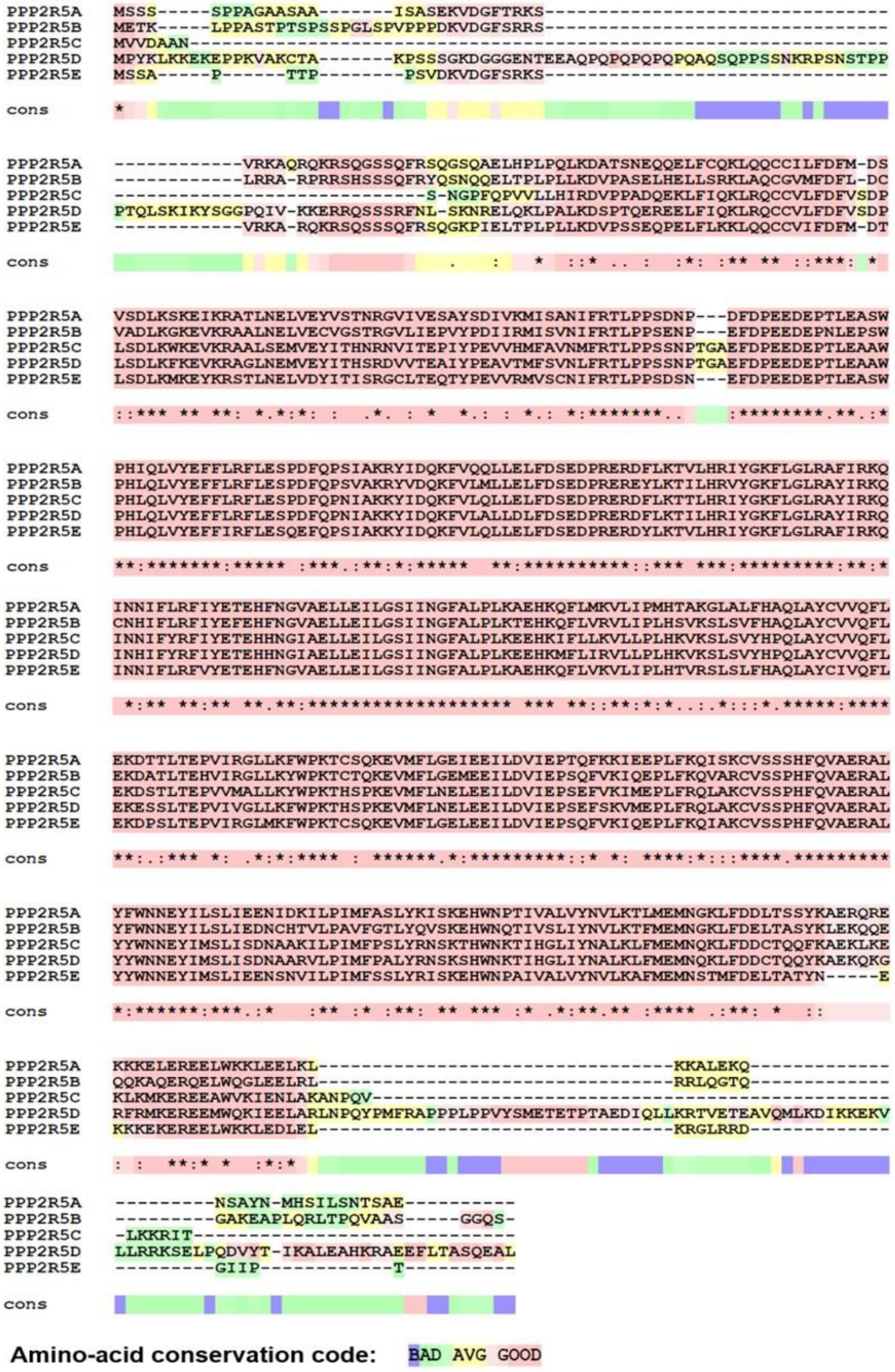


**Extended Data Figure 2. Sequence alignment of the B56 family of PP2A regulatory subunits.** The levels of sequence conservation are color-coded, as shown at the bottom. The alignment was performed using the T-Coffee multiple sequence alignment program.

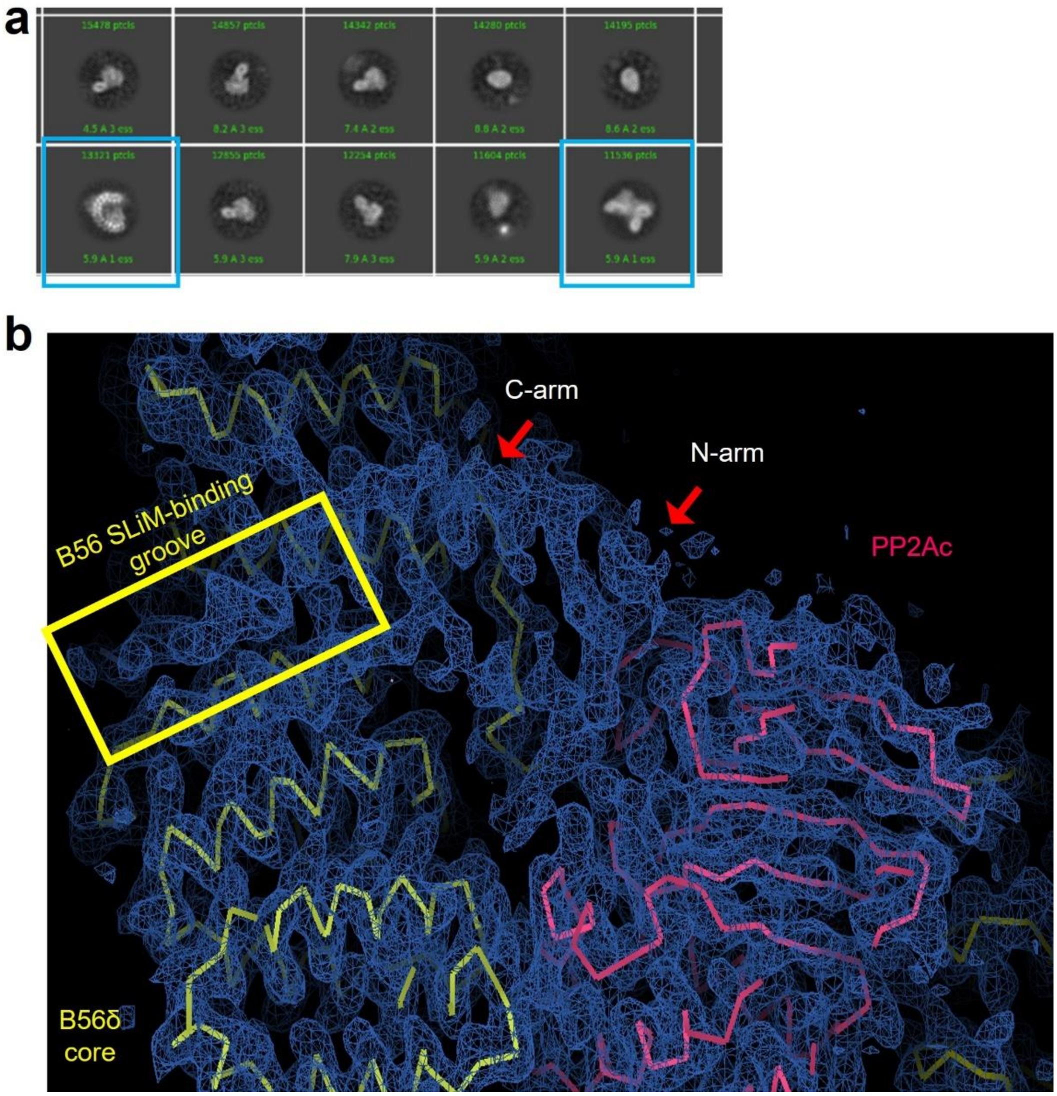


**Extended Data Figure 3. Single-particle cryo-EM of the WT PP2A-B56δ holoenzyme on the Spotiton grids.** (a) Representative 2D class averages gave two classes (cyan boxes) for the intact holoenzyme. Most particles are dissociated subunits with significantly smaller sizes than the intact holoenzyme. (b) The Cryo-EM map resulting from data in (a) fits with the model of the holoenzyme core. It reveals the electron density for the N/C-arms that occupy the B56 SLiM-binding groove (highlighted in yellow rectangular).

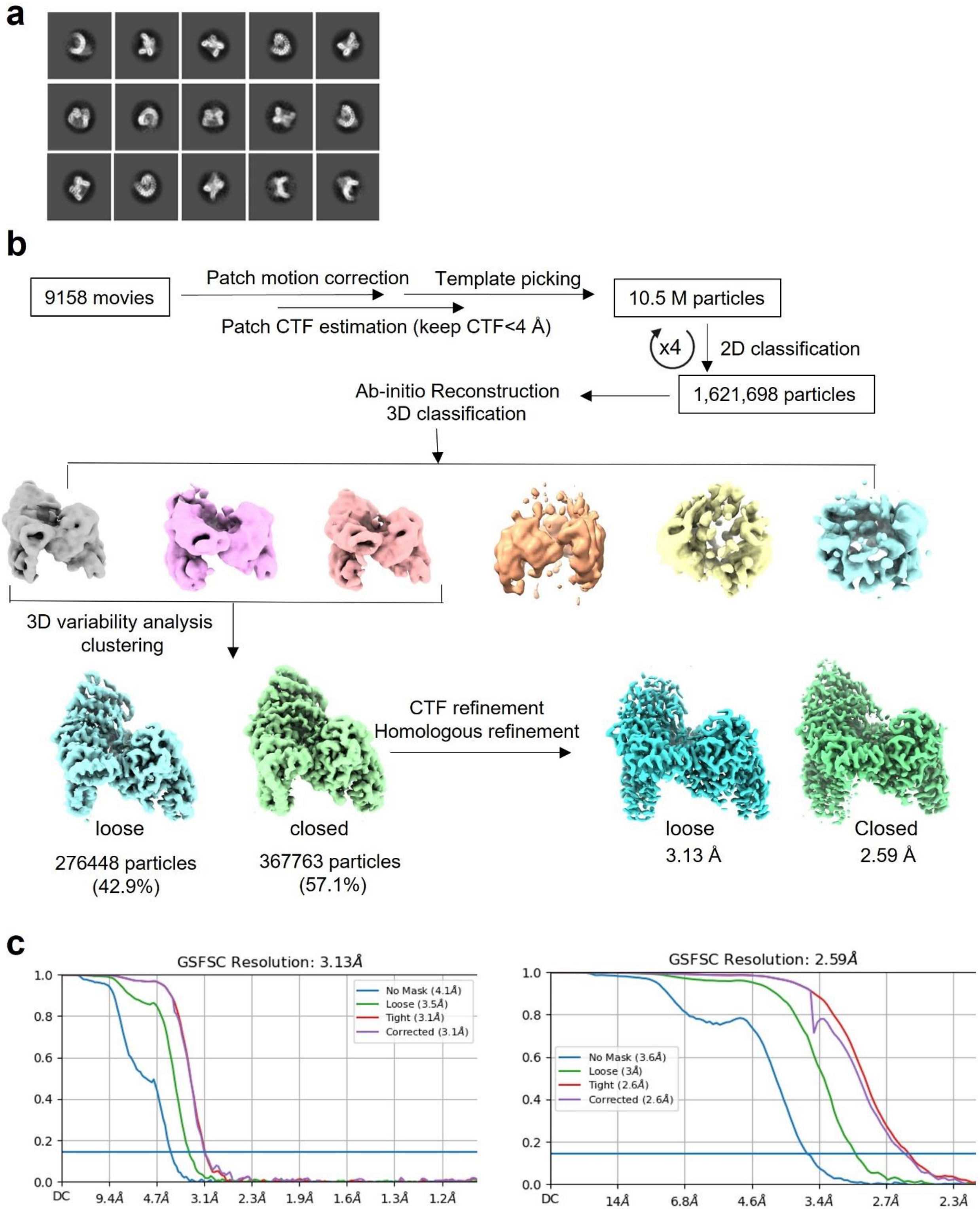


**Extended Data Figure 4. Cryo-EM data processing and reconstruction of the E197K PP2A-B56δ holoenzyme.** (a) The representative 2D class averages of the E197K PP2A-B56δ holoenzyme. (b) Flowchart of the data processing workflow using CryoSPARC[3]. (c) The global resolutions of the closed and open forms of the E197K PP2A-B56δ holoenzyme are 3.13 Å and 2.59Å, respectively, at 0.143 Fourier shell correlation (FSC) as calculated by CryoSPARC.

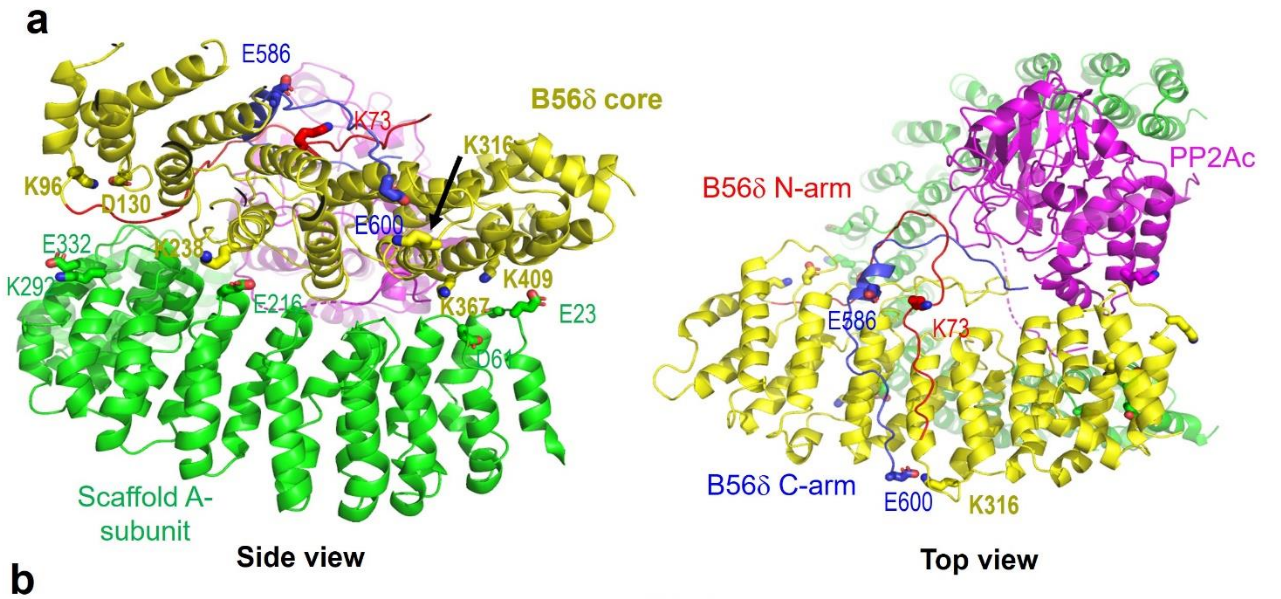


**b**

| Crosslink | Number of experiments detecting the crosslink |
|---|---|
| A-E292/A-K323 | 4 (previous structure) |
| B56δ-D130/B56δ-K96 | 1 (previous structure) |
| A-E216/B56δ-K238 | 2 (previous structure) |
| A-D61/B56δ-K367 | 1 (previous structure) |
| A-E23/B56δ-K409 | 3 (previous structure) |
| B56δ-C-E600/B56δ-K316 | 2 (new interaction) |
| B56δ-C-E586/B56δ-N-K73 | 3 (new interaction) |

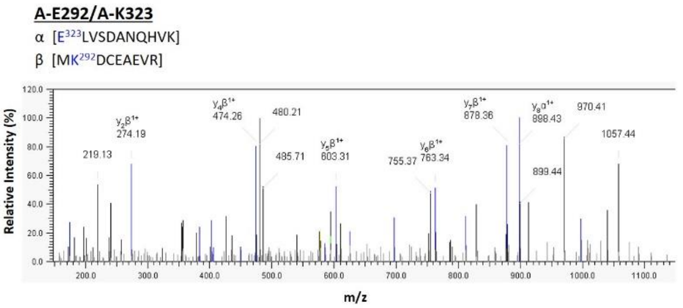


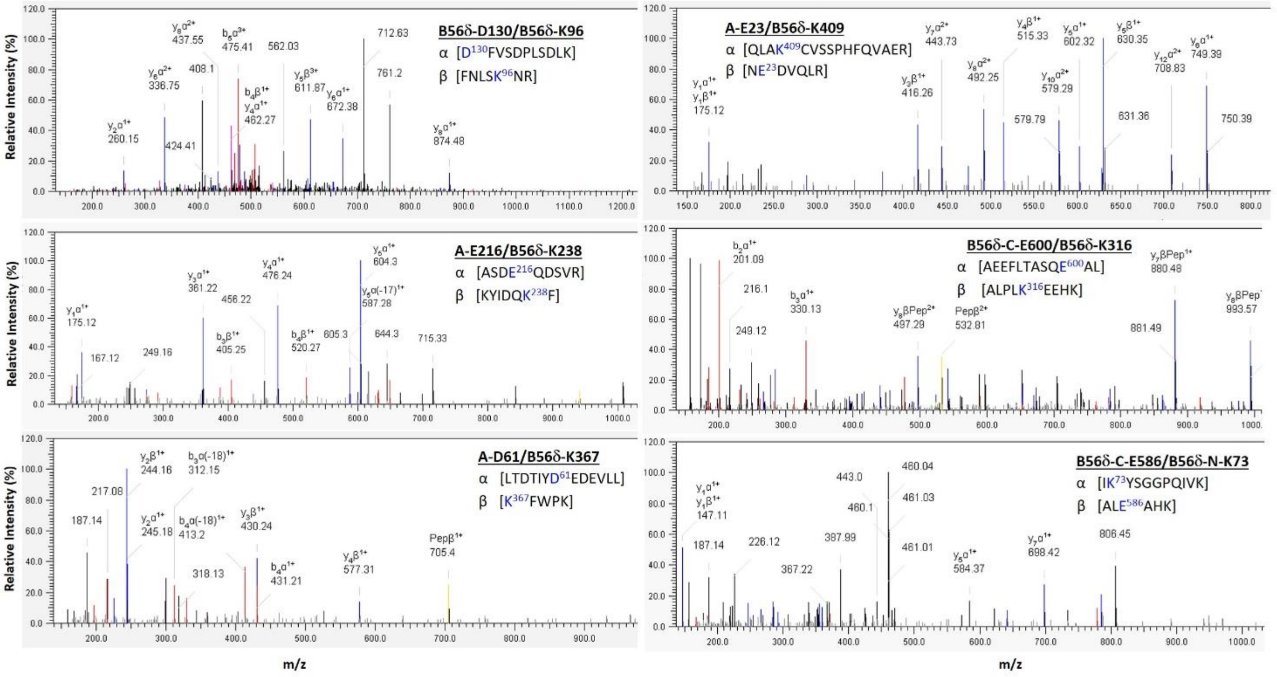


**Extended Data Figure 5. Crosslinking mass spectrometry (XL-MS) identified the intra- and inter-subunit interactions of the PP2A-B56δ holoenzyme.** (a) The side (upper left) and top (upper right) views of the structural model for the closed form of the PP2A-B56δ holoenzyme illustrate the crosslinked residue pairs with their side chains shown in sticks. (b) The table in the upper-left panel summarizes the crosslinked pairs and the frequency of successful detections in four XL-MS experiments. The panels in the upper right and bottom show the MS2 spectra of identified crosslinks. The crosslinked peptides are labeled as peptides α and β. The specific residues that were crosslinked within each peptide are highlighted in blue.

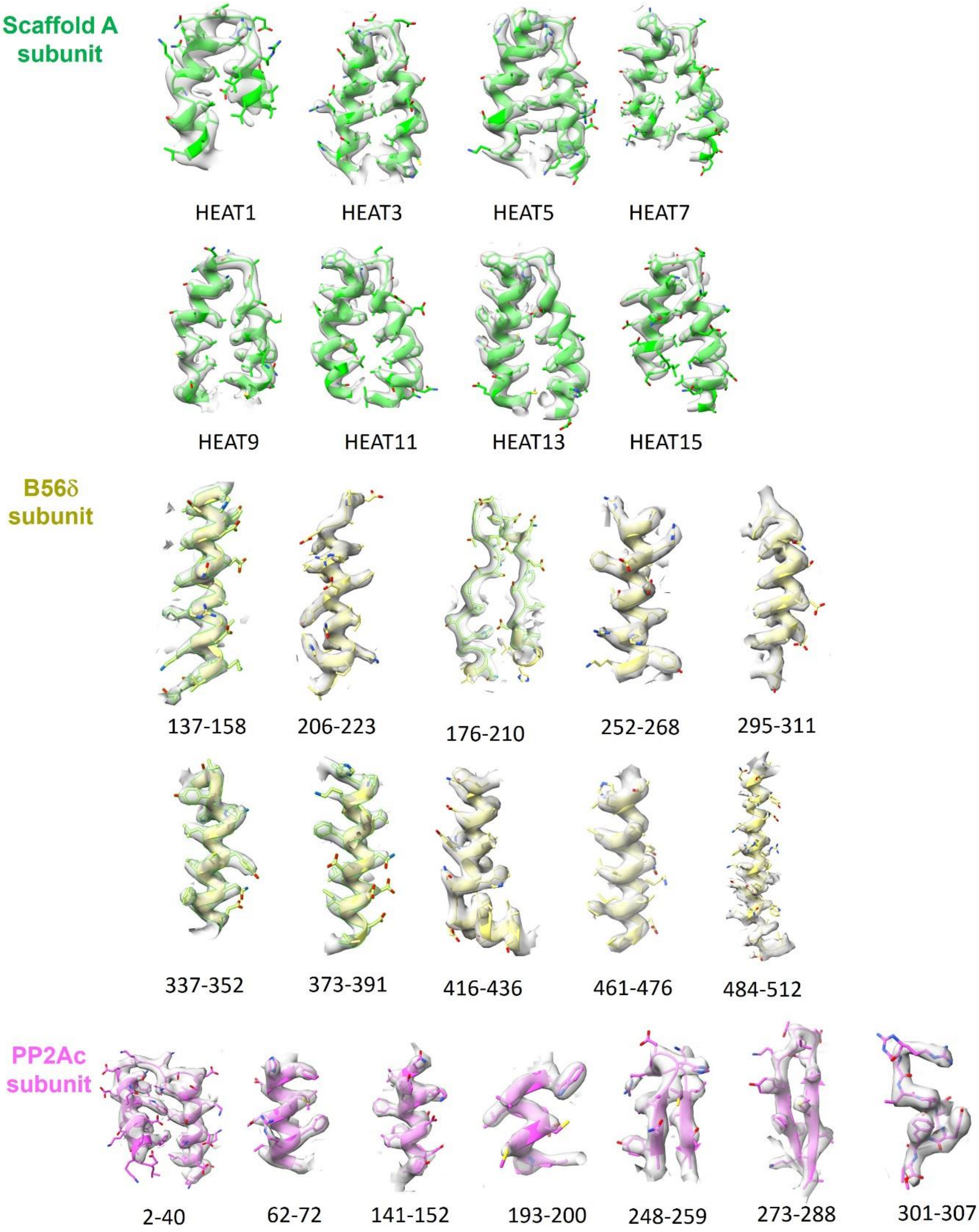


**Extended Data Figure 6.** Electron microscopy densities and models of the scaffold (green), catalytic (magenta), and B56δ (yellow) subunits of E197K PP2A-B56δ holoenzyme showed the agreement between the electron densities and the protein models in the representative regions.

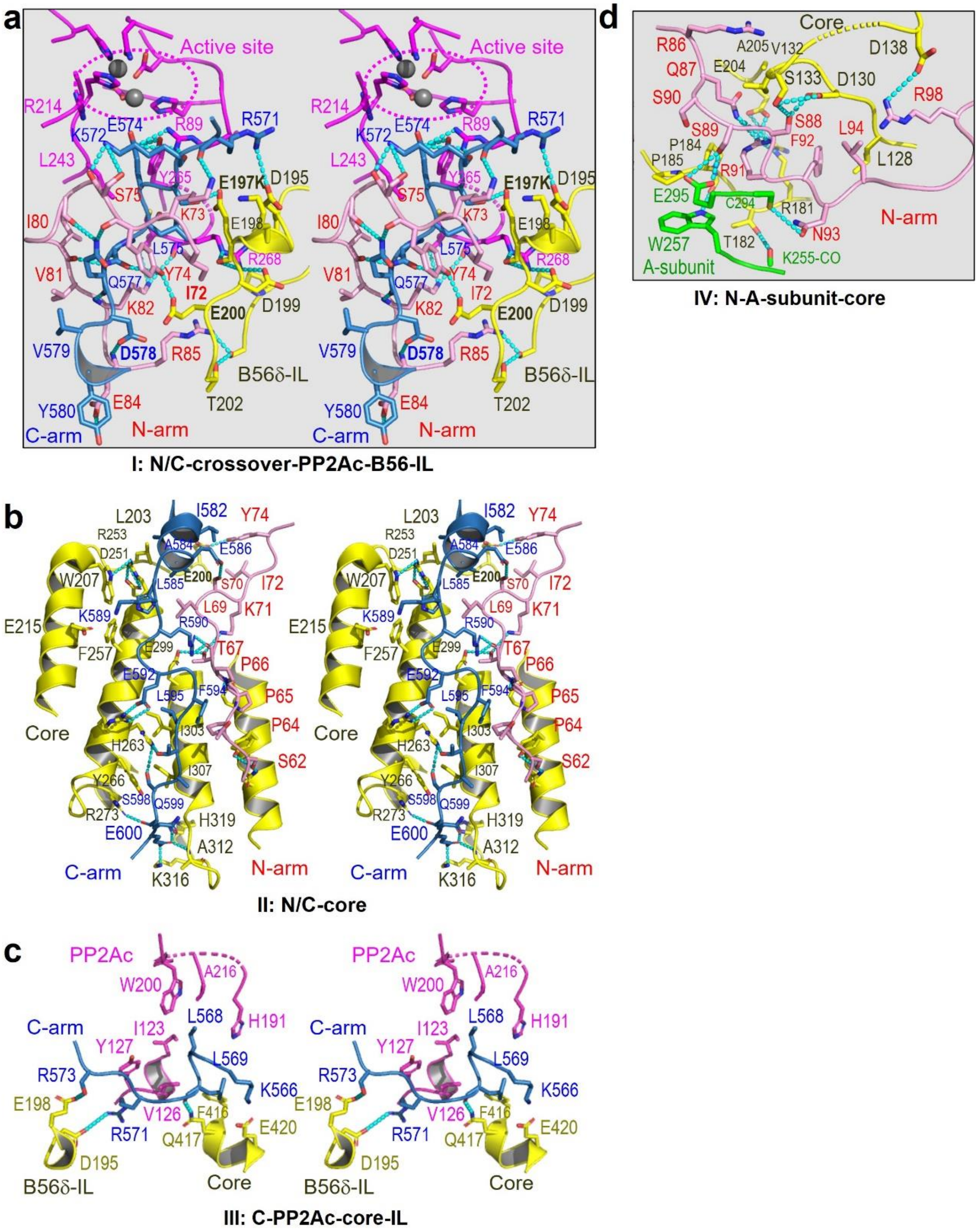


**Extended Data Figure 7. The close-up stereo views of the super-long regulation interface in the PP2A-B56δ holoenzyme.** (a-d) showed four regions of the extended interface divided as in Fig. 1b. The A subunit, PP2Ac, B56δ core, and N/C-arms are shown in cartoon and colored green, magenta, yellow, pink, and blue, respectively. Residues at the interfaces are shown in sticks.

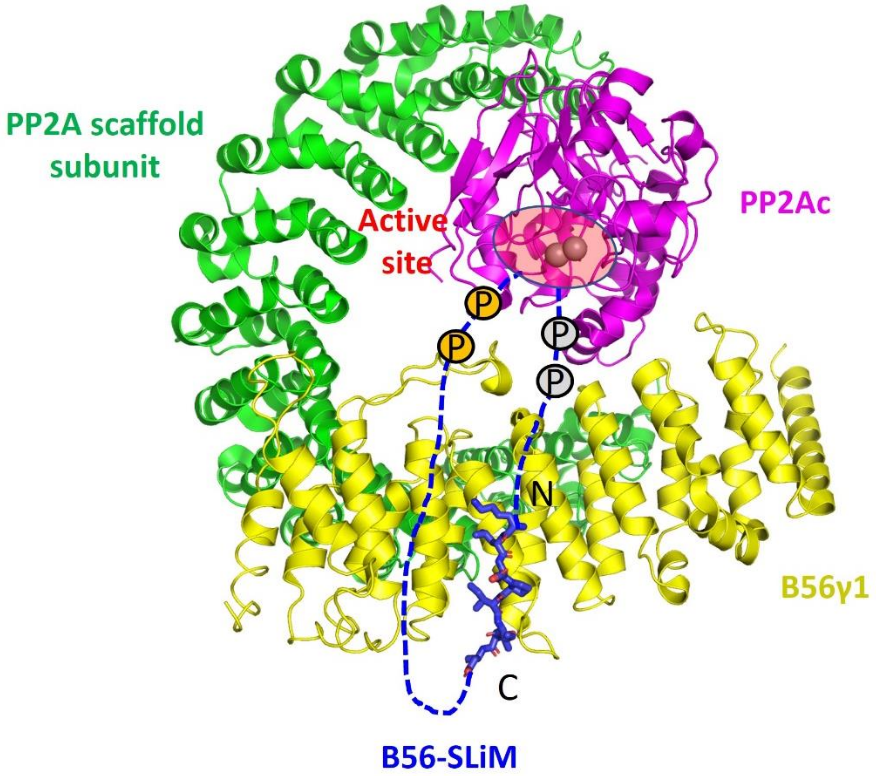


**Extended Data Figure 8. Structural illustration of the B56 SLiM-binding site and the phosphatase active site (red circle) of the PP2A-B56γ1 holoenzyme, representative of the holoenzyme core in the B56 family**. The structure of a B56 SLiM (blue) from BubR1 bound to B′γ1 (yellow; PDB code: 5jja) overlapped with the structure of the holoenzyme (PDB code: 2NYL). The dashed lines and circles stand for peptide fragments and phosphorylation sites upstream (grey) and downstream (orange) of the SLiM. N and C stand for N- and C-termini of the bound SLiM. Manganese ions in the active site are shown in grey spheres.

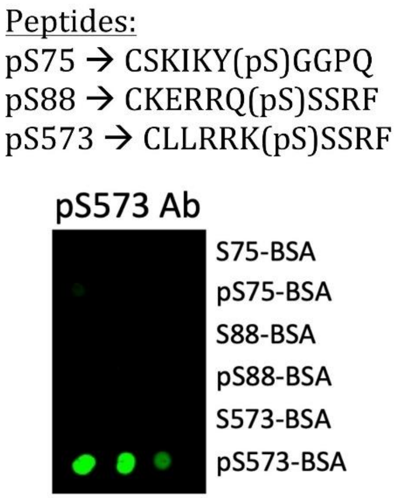


**Extended Data Figure 9. Dot blot analysis of the specificity of pS573 antibody.** The sequences of the phosphorylated (p) peptides covering different regions of the B56δ subunit used for testing are shown above.

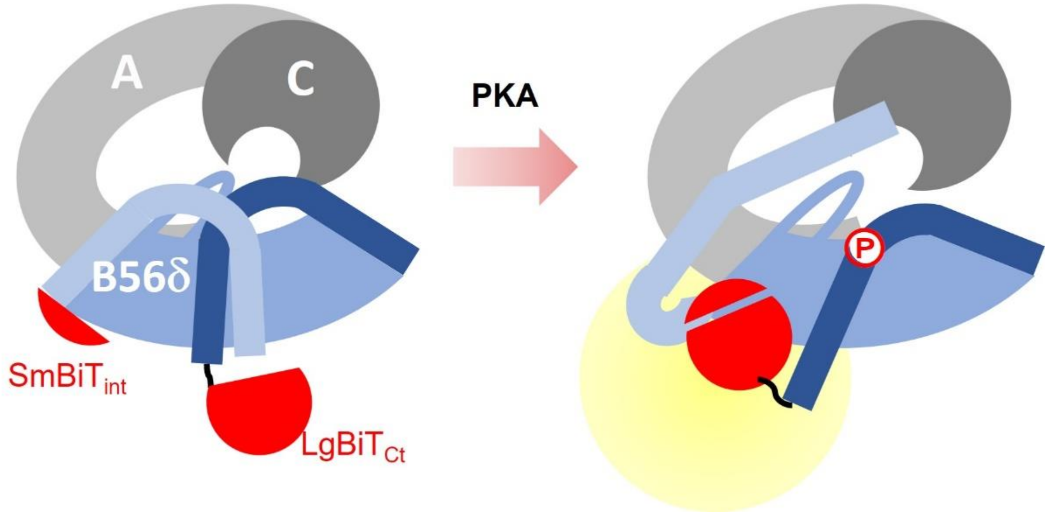


**Extended Data Figure 10. Cartoon illustration of the split NanoBiT B56δ holoenzyme sensor.** The SmBiT peptide fragment was inserted in the holoenzyme core immediately downstream of the N-arm, and the LgBiT fragment was fused to the C-terminus of the C-arm. The two NanoBiT fragments are restricted in separate positions in the closed form of the PP2A-B56δ holoenzyme. Upon activation phosphorylation by PKA, the N/C-arms open up and allow the two fragments to interact and form an active NanoBiT enzyme.

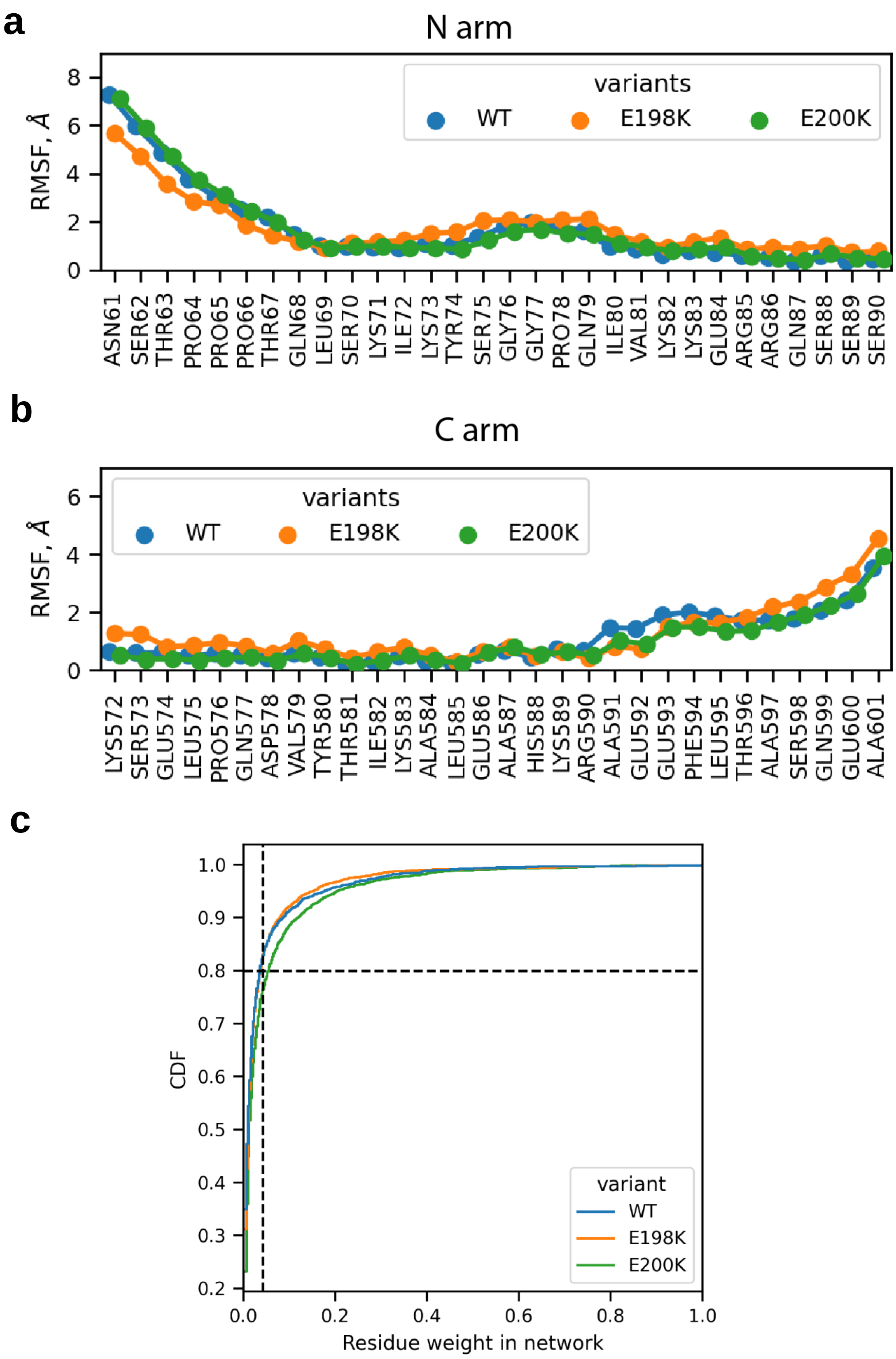


**Extended Data Figure 11.** (a, b) Fluctuation of the N/C arms in REST2 simulations. (c) Cumulative distribution function of residues weights in the allosteric networks of the PP2A holoenzyme. The selected cutoff at 80$^{th}$ percentile is shown with dashed lines and corresponds to residue weight 0.042.

**Extended Data Table 1. Cryo-EM data collection, refinement and validation statistics**

| PP2A-B56δ holoenzyme | WT closed form by Spoiton | E197K closed form |
|---|---|---|
| Data collection and processing | | |
| Magnification (kx) | 105 | 81 |
| Voltage (kV) | 300 | 300 |
| Defocus (μm) | -1.2 to -2 | -0.7 to -2.2 |
| Pixel size (Å) | 1.096 | 1.068 |
| Total dose ($e^-$ / $Å^2$) | 66.84 | 49 |
| Number of frames | 50 | 69 |
| Number of micrographs | 1790 | 9158 |
| Number of micrographs used | 1534 | 7305 |
| Number of initial particles picked | 1,845,319 | 10,502,924 |
| Number of final particles refined | 40778 | 367,763 |
| Map resolution (Å) | 4.00 | 2.59 |
| FSC threshold (Å) | 0.143 | 0.143 |
| Refinement | | |
| Model cutoff (Å) | 4.0 | 2.7 |
| Total number of atoms | 10136 | 11036 |
| Map sharpening B factor ($Å^2$) | 73.1 | 97.9 |
| RMSD | | |
| bond length ($Å^2$) | 0.007 | 0.005 |
| bond angle (°) | 0.897 | 0.610 |
| CC | 0.79 | 0.85 |
| Validation | | |
| MolProbity Score | 2.55 | 1.60 |
| Clash score | 33.70 | 6.88 |
| Ramachandran plot (%) | | |
| Favored | 90.33 | 96.56 |
| Allowed | 9.68 | 3.44 |
| Disallowed | 0 | 0 |
